# Statistics or biology: the zero-inflation controversy about scRNA-seq data

**DOI:** 10.1101/2020.12.28.424633

**Authors:** Ruochen Jiang, Tianyi Sun, Dongyuan Song, Jingyi Jessica Li

**Author notes:** Twitter handles: Ruochen Jiang (@Ruochen_Jiang); Tianyi Sun (@TianyiSun17); Dongyuan Song (@SongDongyuan); Jingyi Jessica Li (@jsb_ucla).

## Abstract

Researchers view vast zeros in single-cell RNA-seq data differently: some regard zeros as biological signals representing no or low gene expression, while others regard zeros as missing data to be corrected. To help address the controversy, here we discuss the sources of biological and non-biological zeros; introduce five mechanisms of adding non-biological zeros in computational benchmarking; evaluate the impacts of non-biological zeros on data analysis; benchmark three input data types: observed counts, imputed counts, and binarized counts; discuss the open questions regarding non-biological zeros; and advocate the importance of transparent analysis.

## Introduction

The rapid development of single-cell technologies has brought unprecedented opportunities to quantifying transcriptome heterogeneity among individual cells and transcriptome dynamics along cell developmental trajectories [1–4]. Many single-cell RNA sequencing (scRNA-seq) protocols have been developed. Two major types of protocols are (1) tag-based, unique molecular identifier (UMI)-based protocols such as Drop-seq [5] and 10x Genomics Chromium [6, 7] and (2) full-length, non-UMI-based protocols such as Smart-seq2 [8] and Fluidigm C1 [9]. Different protocols exhibit disparate accuracy and noise levels for quantifying gene expression in single cells, posing many computational and analytical challenges for researchers to extract biological knowledge from scRNA-seq data [10, 11]. Facing these challenges, computational researchers have developed hundreds of computational and statistical methods for various scRNA-seq data analytical tasks, including the selection of informative marker genes [12–16], the identification of cell types and states [14, 17–23], the reconstruction of cell developmental trajectories [24–29], and the identification of cell-type-specific genes [13, 28, 30–38].

A universal analytical challenge for scRNA-seq data generated by any protocol is the vastly high proportion of genes with zero expression measurements in each cell. This data sparsity issue is apparent when scRNA-seq data are compared with bulk RNA-seq data [36, 39, 40], which contain aggregated gene expression measurements from many cells. While the proportion of zeros in bulk RNA-seq data is usually 10%–40% [41, 42], that proportion can be as high as 90% in scRNA-seq data [43]. Such excess zeros would bias the estimation of gene expression correlations [44] and hinder the capture of gene expression dynamics [45] from scRNA-seq data. In early scRNA-seq data analyses, the high data sparsity provoked the use of zero-inflated models [36, 38, 46] and the development of imputation methods for reducing zeros [20, 44, 45, 47–63]. More recently, however, there were voices against the use of zero-inflated models for scRNA-seq data generated by UMI protocols [64, 65]. Besides, there was a proposal for treating zeros as useful information that researchers should embrace [66]. These mixed statements raised a fundamental question to the scRNA-seq field: should we use or remove zeros in scRNA-seq data analysis?

In this article, we provide some perspectives on this puzzling question by discussing the sources of zeros in scRNA-seq data, the impacts of zeros on various data analyses, the existing approaches for handling zeros, and the pros and cons of these approaches. Specifically, first, we define biological and non-biological zeros arising from scRNA-seq data generation, and we clarify several ambiguous terms about zeros in the scRNA-seq literature. Second, we use scRNA-seq data generated by Drop-seq, 10x Genomics, and Smart-seq2 to demonstrate the relation between zero patterns and protocols. Further, we use simulation studies to evaluate the effects of zeros and zero-generation mechanisms on cell clustering and differentially expressed (DE) gene identification. Third, we summarize three commonly used approaches for handling zeros— direct statistical modeling, imputation, and binarization—and discuss their respective pros and cons. Fourth, we benchmark the performance of the three input data types in three downstream analyses: cell clustering, cell dimension reduction, and DE gene identification. Last, we provide practical advice and outline future directions for bioinformatics tool developers and users.

Table 1 summarizes the key concepts used in this paper, including their definitions and categories (biology, technology, and modeling). Table 2 clarifies three zero-related terms: dropouts, excess zeros, and zero inflation; the first two have been ambiguously used in the literature.

**Table 1:**
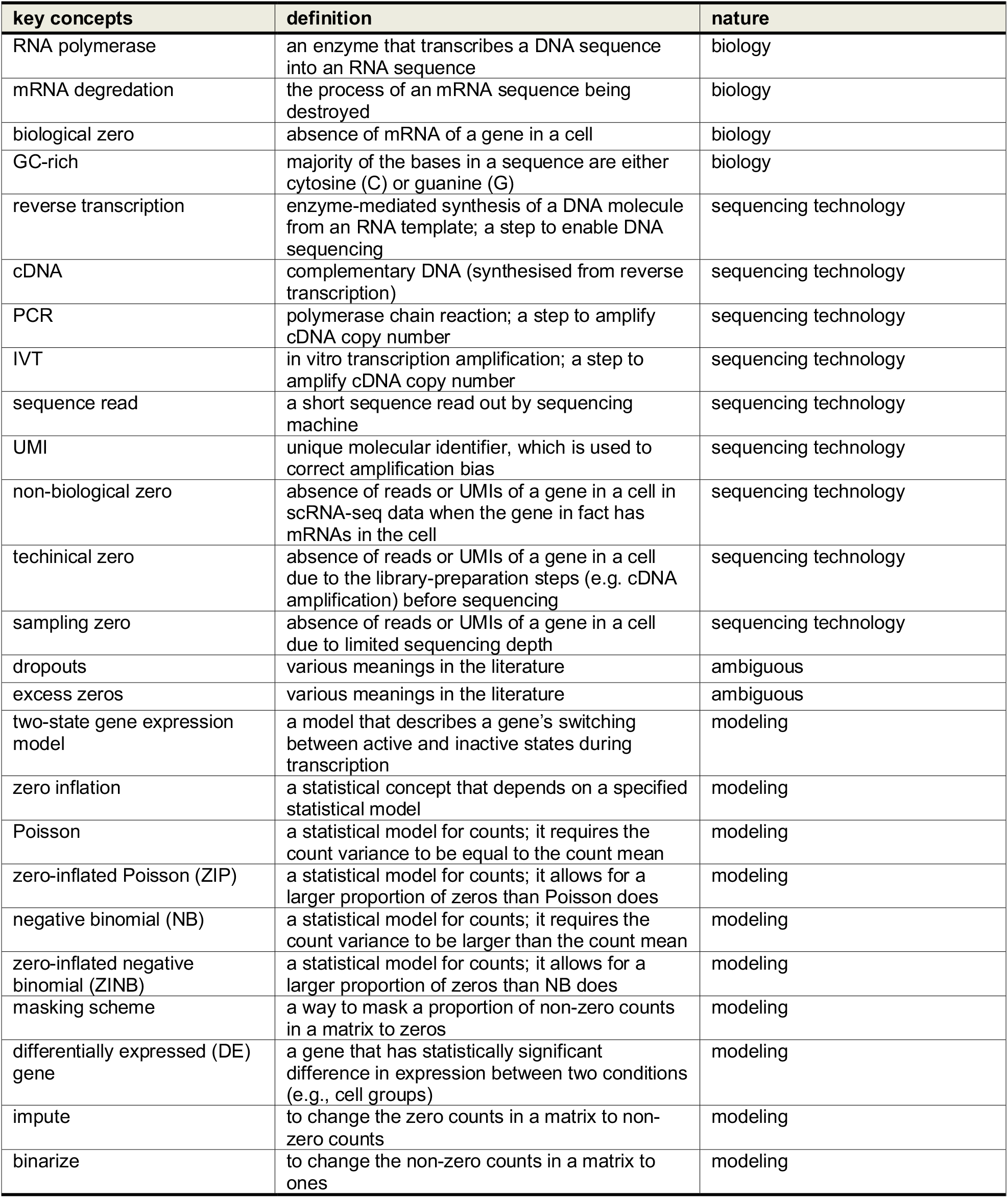
A summary of the key concepts used in this paper, including their definitions and nature.

**Table 2:**
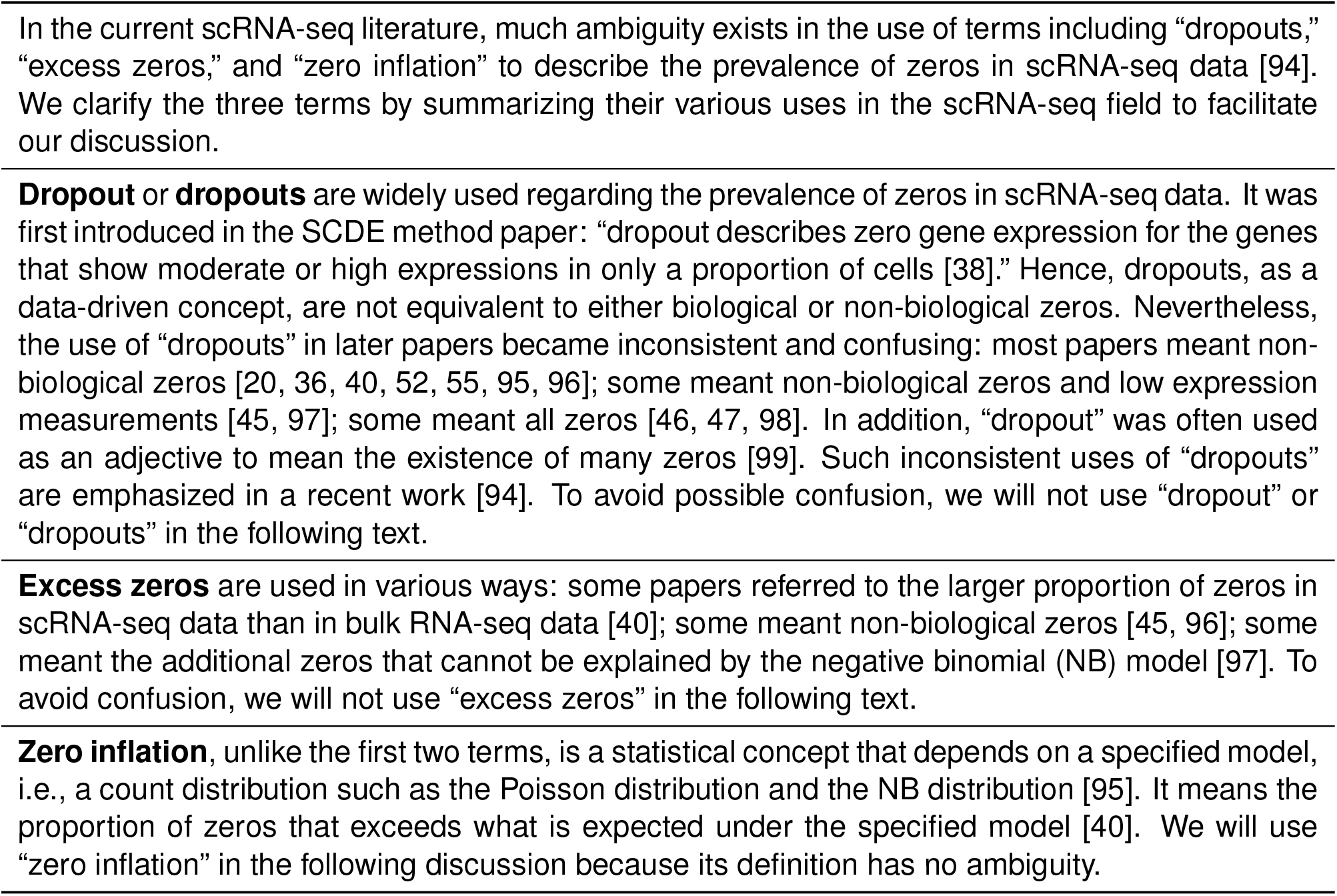
Clarification of zero-related terminology

### Sources of zeros in scRNA-seq data

Zero measurements in scRNA-seq data have two sources: biological and non-biological (Fig. 1). While biological zeros carry meaningful information about cell states, non-biological zeros represent missing values artificially introduced during the generation of scRNA-seq data. In our paper, non-biological zeros include technical zeros, which occur during the preparation of biological samples for sequencing, and sampling zeros, which arise due to limited sequencing depths. Our classification of zeros in sequencing data into biological, technical, and sampling zeros is aligned with the classification in Silverman et al. [67] except a slight difference (we refer to the zeros due to inefficient amplificaiton, e.g., PCR, as sampling zeros, while Silverman et al called them technical zeros). The non-biological zeros have typically been viewed as impediments to the full and accurate interpretation of cell states and the differences between them. Fig. 1a provides an overview of a scRNA-seq experiment, and it highlights the biological factors and technical procedures that may lead to zeros in scRNA-seq data. Fig. 1b summarizes how biological factors result in biological zeros and how technical procedures cause non-biological zeros, including technical zeros and sampling zeros. It is worth noting that biological and non-biological zeros are hardly distinguishable in scRNA-seq data without biological knowledge or spike-in control (see Future directions).

**Figure 1:**
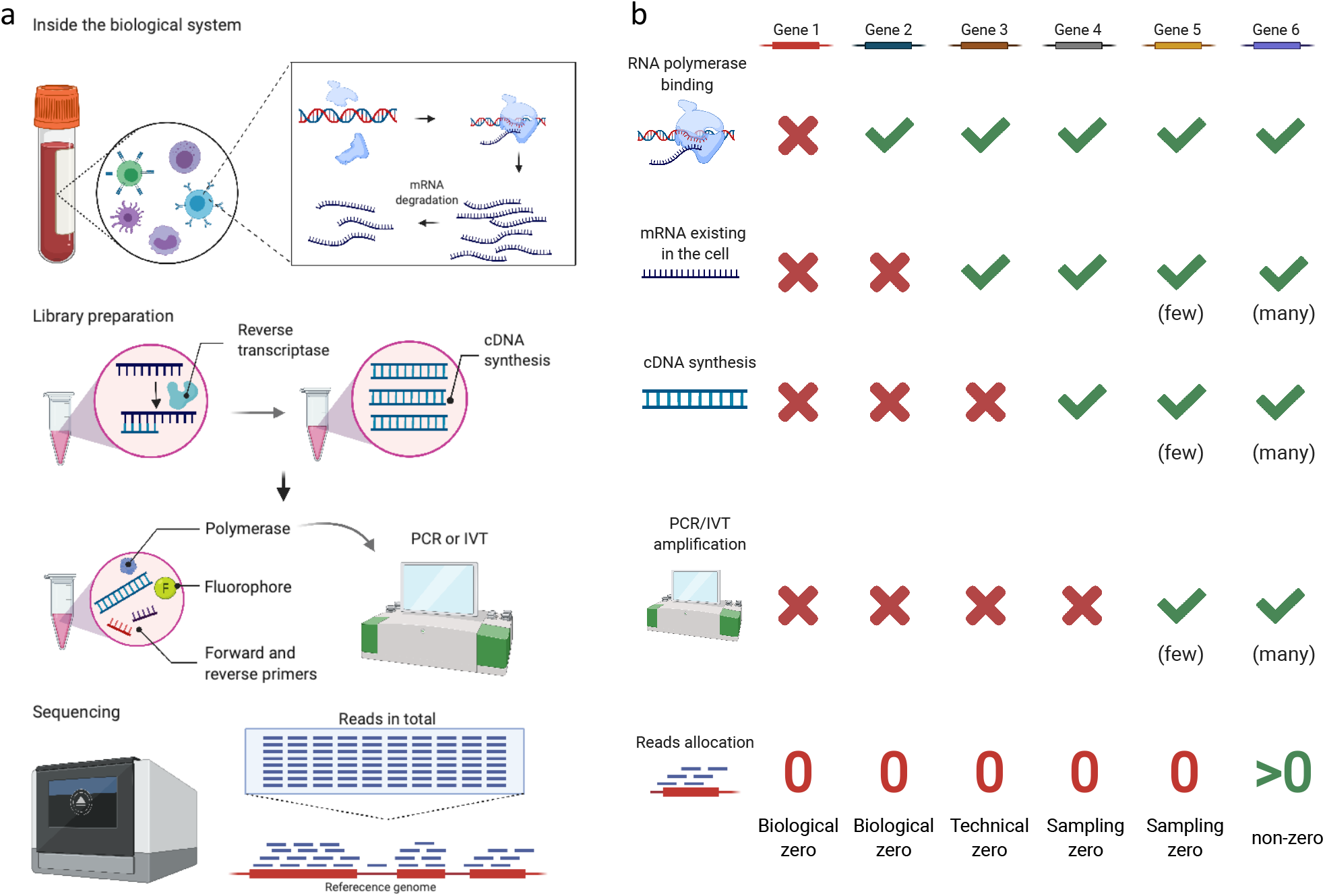
Sources of zeros in scRNA-seq data. **(a)** An overview of a scRNA-seq experiment. Biological factors that determine true gene expression levels include transcription and mRNA degradation (top panel). Technical procedures that affect gene expression measurements include cDNA synthesis, PCR or IVT amplification, and sequencing depth (bottom three panels). Finally, every gene’s expression measurement in each cell is defined as the number of reads mapped to that gene in that cell. **(b)** How the biological factors and technical procedures in (a) lead to biological, technical, and sampling zeros in scRNA-seq data. Red crosses indicate occurrences of zeros, while green checkmarks indicate otherwise. Biological zeros arise from two scenarios: no transcription (gene 1) or no mRNA due to faster mRNA degradation than transcription (gene 2). If a gene has mRNAs in a cell, but its mRNAs are not captured by cDNA synthesis, the gene’s zero expression measurement is called a technical zero (gene 3). If a gene has cDNAs in the sequencing library, but its cDNAs are too few to be captured by sequencing, the gene’s zero expression measurement is called a sampling zero. Sampling zeros occur for two reasons: a gene’s cDNAs have few copies because they are not amplified by PCR or IVT (gene 4), or a gene’s mRNA copy number is too small so that its cDNAs still have few copies after amplification (gene 5). If the factors and procedures above do not result in few cDNAs of a gene in the sequencing library, the gene would have a non-zero measurement (gene 6). The figure is created with BioRender.com.

### Biological zeros in scRNA-seq data

A biological zero is defined as the true absence of a gene’s transcripts or messenger RNAs (mRNAs) in a cell [67]. Biological zeros occur for two reasons (Fig. 1b). First, many genes are unexpressed in a cell (e.g., gene 1 in Fig. 1b), and cells of distinct types have different genes expressed—a fact that results in the diversity of cell types [68, 69]. Second, many genes undergo a bursty process of transcription (i.e., mRNA synthesis); that is, these genes are not transcribed constantly but intermittently, a well-known phenomenon in gene regulation [38, 39, 46, 70–72]. Specifically, in eukaryotic cells, transcription is initiated by the binding of specific transcription factors (TFs) and RNA polymerase to the promoter of a gene [73–75]. Due to the stochasticity of TF binding, a gene switches between active and inactive states, and its transcription only occurs during the active state [76]. Hence, systems biologists have used a two-state gene expression model to describe how the rates of three processes—active/inactive state switching, transcription, and mRNA degradation—jointly determine the distribution of a gene’s mRNA copy numbers, i.e., expression levels, in cells of the same type [76–78]. Fig. 2 illustrates the model and provides three example settings of model parameters along with their corresponding gene expression distributions. Depending on the gene’s switching rates between the active and inactive states, transcription rate, and degradation rate, the resulting distribution may exhibit a mode near zero, which makes it appear that the gene expresses no mRNA at a particular time, in a large number of cells (e.g., gene 2 in Fig. 1b).

**Figure 2:**
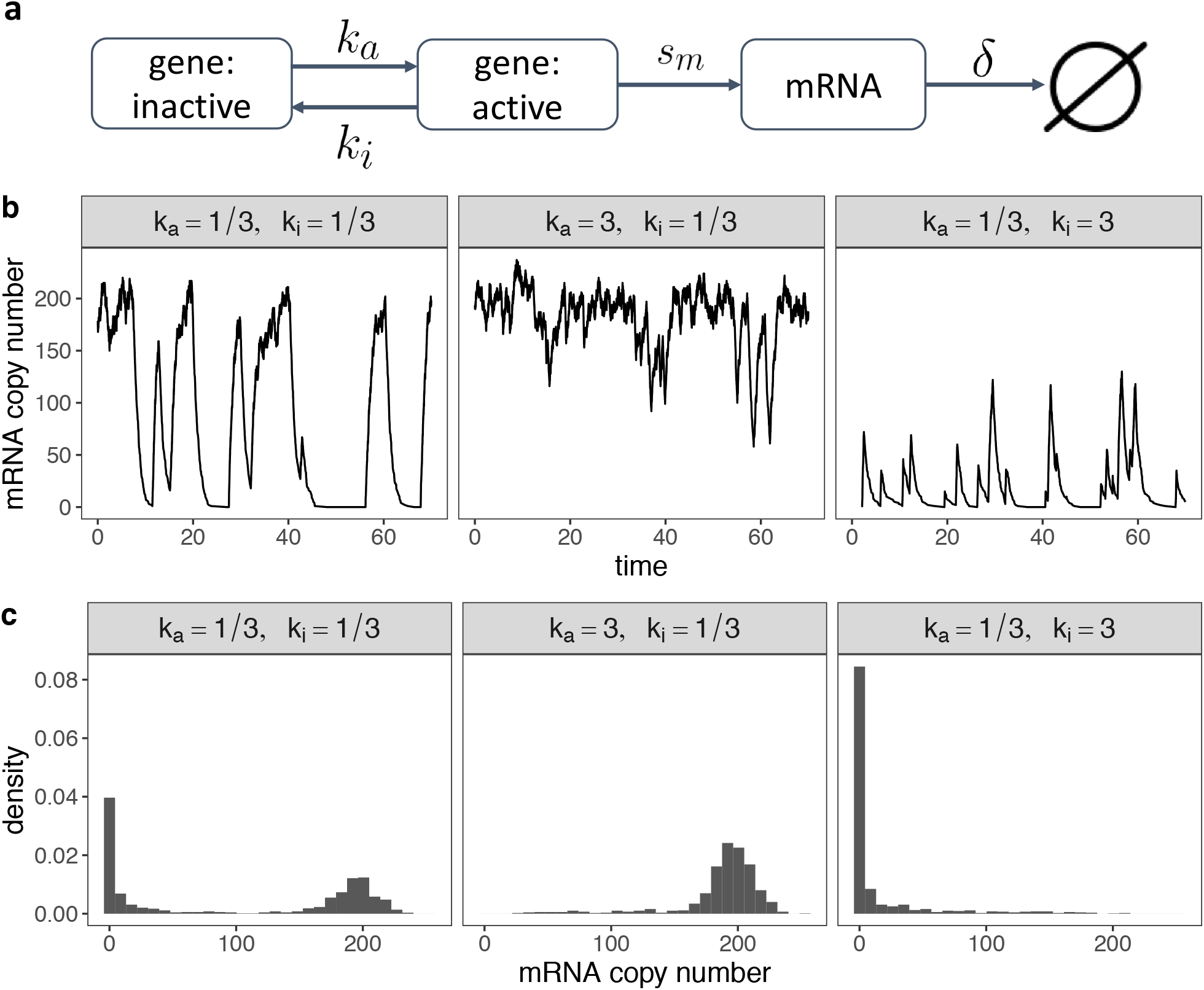
A two-state stochastic model of the expression levels of one gene. **(a)** A diagram of the two-state gene expression model [76–78], where a gene stochastically switches from an inactive state to an active state at rate *k*_*a*_ and from an active state to an inactive state at rate *k*_*i*_. The gene transcribes mRNA at rate *s*_*m*_ only when it is in the active state. The transcribed mRNA then degrades at rate *δ*. **(b)** Given *s*_*m*_ = 200 and *δ* = 1, the effects of *k*_*a*_ and *k*_*i*_ on the temporal dynamics of the gene’s mRNA copy number. Three example values of *k*_*a*_ and *k*_*i*_ are provided. Left: when both *k*_*a*_ and *k*_*i*_ are small, the mRNA copy number switches between small and large values. Middle: when *k*_*a*_ is much larger than *k*_*i*_, the mRNA copy number remains large most of the time. Right: when *k*_*a*_ is much smaller than *k*_*i*_, the mRNA copy number remains small most of the time. **(c)** Distributions of the gene’s mRNA copy number (across cells) corresponding to the three example settings in (b). Left: when the gene’s mRNA copy number switches between small and large values, the resulting distribution is bimodal with two modes at zero and around *s*_*m*_*/δ*. Middle: when the gene’s mRNA copy number is large most of the time, the resulting distribution has a single mode around *s*_*m*_*/δ*. Right: when the gene’s mRNA copy number is small most of the time, the resulting distribution has a single mode at zero. In summary, when *k*_*a*_ is small, the gene is expected to have biological zeros in cells with non-negligible probability.

### Non-biological zeros in scRNA-seq data

Non-biological zeros reflect the loss of information about truly expressed genes due to the inefficiencies of the technologies employed from sample collection to sequencing. Unlike biological zeros, non-biological zeros refer to the zero expression measurements of genes with transcripts in a cell. There are two types of non-biological zeros [67]: (1) technical zeros, which arise from library-preparation steps before sequencing, and (2) sampling zeros, which result from a limited sequencing depth.

One cause of technical zeros is the imperfect mRNA capture efficiency in the reverse transcription (RT) step from mRNA to cDNA. The efficiency has a considerable variation across protocols and may be as low as 20% [79], depending on multiple experimental parameters [80]. The efficiency may even differ between mRNA transcripts. For example, if an mRNA transcript has an intricate secondary structure or is bound to proteins, it would not be reversely transcribed to cDNA efficiently [10, 33, 38]. In summary, if a gene’s mRNA transcripts in a cell are not converted into cDNA molecules (cDNAs), the gene would falsely appear as non-expressed in that cell in the sequencing library, resulting in a technical zero in scRNA-seq data (e.g., gene 3 in Fig. 1b).

The other type of non-biological zeros, sampling zeros, occurs due to a constraint on the total number of reads sequenced, i.e., the sequencing depth [64, 81], which is determined by the experimental budget and sequencing machine. During sequencing, cDNAs are randomly captured (“sampled”) and sequenced into reads. Hence, a gene with fewer cDNAs is more likely to be undetected due to this random sampling. If undetected, the gene’s resulting zero read count is a “sampling zero.” There are two reasons why a gene (in a cell) may have too few cDNAs in the sequencing library: too few cDNAs before amplification and inefficient cDNA amplification. Below we explain why cDNA amplification may cause some genes to have a disproportionally low cDNA copy number in the sequencing library (see Fig. 3).

**Figure 3:**
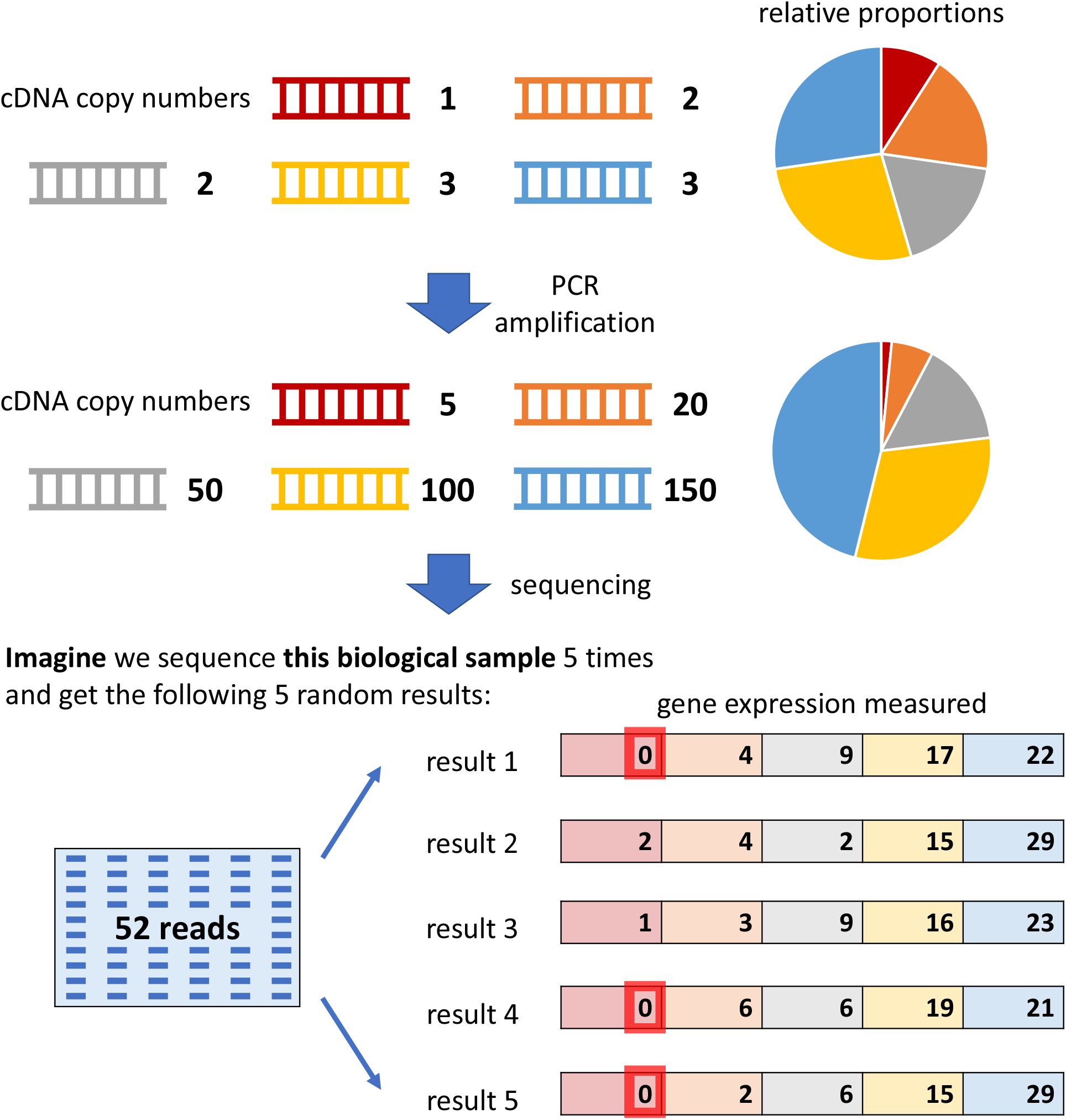
A toy example showing how the PCR amplification may result in sampling zeros. Five genes have their cDNAs amplified by PCR. After the non-linear amplification, their relative proportions change. If the sequencing depth is limited to 52 reads, the first gene has sampling zeros in three out of five hypothetical sequencing experiments.

The cDNA amplification step is essential for scRNA-seq, as it increases the number of cDNA copies of a gene so that the gene is more likely to be detected by sequencing. Polymerase chain reaction (PCR) [82] is the most widely-used amplification procedure. However, PCR amplification is non-linear; thus, the ratio between the copy numbers of two differentially expressed genes is artificially distorted by PCR, i.e., a ratio greater (or smaller) than one becomes even larger (or smaller) after PCR. As a remedy, *in vitro* transcription (IVT) has been developed for linear amplification [83]. However, compared with PCR, IVT requires more input cDNAs to ensure successful amplification; thus, PCR is still the dominant amplification procedure for scRNA-seq [84]. Though indispensable, cDNA amplification is known to introduce biases into cDNA copy numbers because the amplification efficiency depends on cDNA sequence and structure [85, 86]. For example, GC-rich cDNA sequences are harder to be amplified [67]. The amplification efficiency also depends on the design of cell barcodes, adapters, and primers; overlaps or complementarity of barcode, adapter, and primer sequences would induce cDNA secondary structures and thus reduce the amplification efficiency [87, 88]. Moreover, cDNA copy number biases would accumulate as the number of amplification cycles increases [85, 89]; that is, the more cycles, the larger the difference of two genes (with different amplification efficiency) in cDNA copy numbers [90, 91]. Since different scRNA-seq experiments may use different numbers of amplification cycles (e.g., 18 cycles in a Smart-seq2 experiment [92] and 14 cycles in a 10x Genomics experiment [93]), cDNA copy number biases differ among scRNA-seq datasets. In addition, the non-linear amplification nature of PCR would exaggerate the expression level differences between lowly-expressed and highly-expressed genes. Altogether, due to amplification biases, cDNA copy numbers in a sequencing library may not reflect cDNAs’ actual proportions before amplification. As a result, the genes with small cDNA proportions in the sequencing library are likely to be missed by sequencing and thus result in sampling zeros (e.g., gene 4 suffering from inefficient amplification and gene 5 having too few cDNAs in Fig. 1b).

### Debate on zero-inflated modeling of scRNA-seq data

Since the advent of scRNA-seq, zero-inflated models have been widely used in bioinformatics tools on the observed scRNA-seq count data [36, 38, 46, 100]. Zero-inflated models are mixture probabilistic models with two components: a point mass at zero and a common distribution, including the Poisson and NB distributions for read or UMI counts and the normal distribution for log-transformed read or UMI counts. More recently, however, researchers have found that UMI counts are not zero-inflated when compared with the Poisson or NB distribution [6, 65, 67, 93, 94]. In particular, Kim *et al*. provided comprehensive evidence that zero UMI counts can be accounted for by either the NB distribution or even simply the Poisson distribution [65].

The use of UMIs in scRNA-seq can correct the amplification biases in non-zero gene expression measurements [101]; that is, UMIs can be used to identify and remove reads from cDNA duplicates that are results of amplification, and thus some non-zero gene expression measurements would be reduced. However, UMIs cannot help recover sampling zeros, whose corresponding cDNA copy numbers stay unknown despite the use of UMIs [100]. In fact, UMIs cannot reduce any zeros, including biological and non-biological ones. The change of modeling choice—from zero-inflated models for non-UMI-based data to non-zero-inflated models for UMI-based data— indicates that whether or not to use zero-inflated models has nothing to do with the prevalence of zeros. In other words, the modeling choice is a statistical consideration and says nothing about the proportions of zeros or the distinction between biological and non-biological zeros.

Four count distributions—Poisson, zero-inflated Poisson (ZIP), NB, and zero-inflated NB (ZINB)— have been widely used to model a single gene’s read or UMI counts across cells in scRNA-seq data. In fact, the former three models are special cases of the ZINB model (Fig. 4a). Poisson only has one parameter (*λ*) equal to both mean and variance (Fig. 4b). Compared with Poisson, ZIP has one more zero-inflation parameter (*p*) to indicate the proportion of additional zeros that do not come from Poisson (Fig. 4c); when this zero-inflation parameter is zero, ZIP reduces to Poisson. Also, compared with Poisson, NB has one more dispersion parameter (*ψ*) that indicates the over-dispersion of variance relative to the mean (i.e., unlike Poisson, NB has variance greater than mean; Fig. 4d); when this dispersion parameter is positive infinity, NB reduces to Poisson. Compared with NB, ZINB has one more zero-inflation parameter (*p*) to indicate the proportion of additional zeros that do not come from NB (Fig. 4e); when this zero-inflation parameter is zero, ZINB reduces to NB.

**Figure 4:**
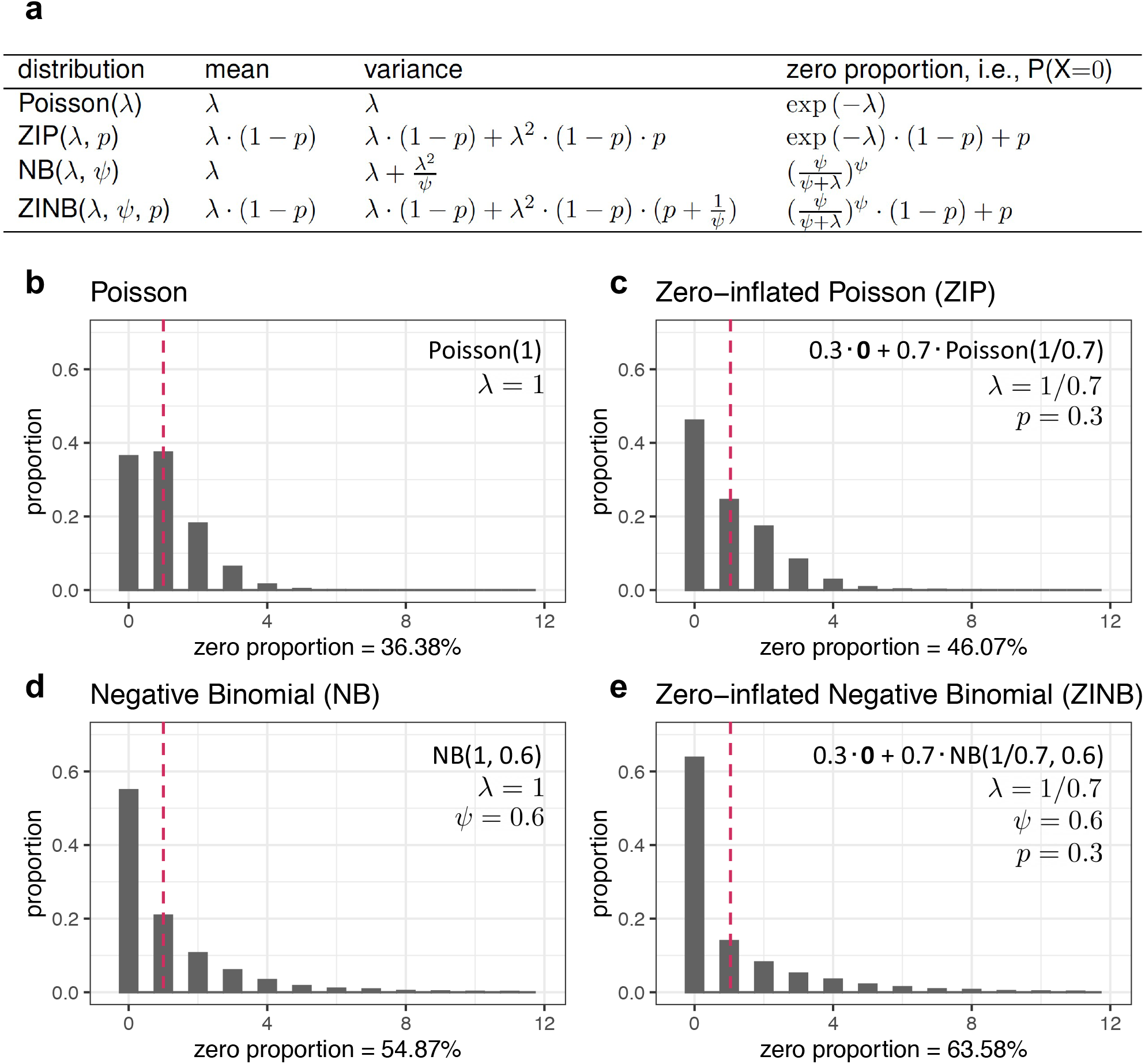
Four count distributions: Poisson, zero-inflated Poisson (ZIP), negative binomial (NB), and zero-inflated negative binomial (ZINB). **(a)** Parameterization, mean, variance, and zero proportion of each of the four distributions. **(b), (c), (d)**, and **(e)** Illustration of the probability mass functions of Poisson (b), ZIP (c), NB (d), and ZINB (e) distributions that all have mean equal to 1. The horizontal axis indicates each possible value, and the vertical axis indicates the probability of taking each possible value. For each distribution, the parameter values are listed on the top right, and the zero proportion is listed at the bottom.

For a fair comparison, we illustrate these four distributions, with example parameters such that they all have the same mean as one (Fig. 4b–e). With the same mean, ZIP and NB have more zeros than Poisson does, and ZINB has the most zeros. Between ZIP and NB, which one has more zeros depends on their parameter values, and when they have the same zero proportion, their non-zero distributions are still different. Moreover, when the four distributions have the same mean, compared with Poisson and ZIP, NB and ZINB have heavier right tails, i.e., greater probabilities of taking larger values.

Svensson shows that non-zero-inflated distributions can describe the variation in droplet scRNA-seq data using droplet-based ERCC spike-in data [64]. To evaluate this claim on real scRNA-seq data with both droplet-based and full-length protocols, we perform the similar analysis on three real scRNA-seq PBMC datasets. More specifically, we fit the above four count distributions—two zero-inflated (ZIP and ZINB) and two non-zero-inflated (Poisson and NB)—to a non-UMI-based dataset generated by Smart-seq2 and two UMI-based datasets generated by 10x Genomics and Drop-seq. These three datasets are ideal for studying how the modeling choice depends on the experimental protocol, as they were generated by a benchmark study [43] that applied multiple scRNA-seq protocols to measure peripheral blood mononuclear cells (PBMCs) from the same batch, and the benchmark study labeled cells using the same cell types and curated genes to be the same across protocols. We first compare the three datasets in terms of their distributions of cell library size (i.e., the total number of reads or UMIs in each cell), numbers of cells, and distributions of the number of genes detected per cell. Fig. 5a–c show that, compared with the two UMI-based datasets, the Smart-seq2 (non-UMI-based) dataset has larger cell library sizes, fewer cells, and more genes detected—a phenomenon consistent across the five cell types (B cells, CD14+ monocytes, CD4+ T cells, cytotoxic T cells, and natural killer cells).

**Figure 5:**
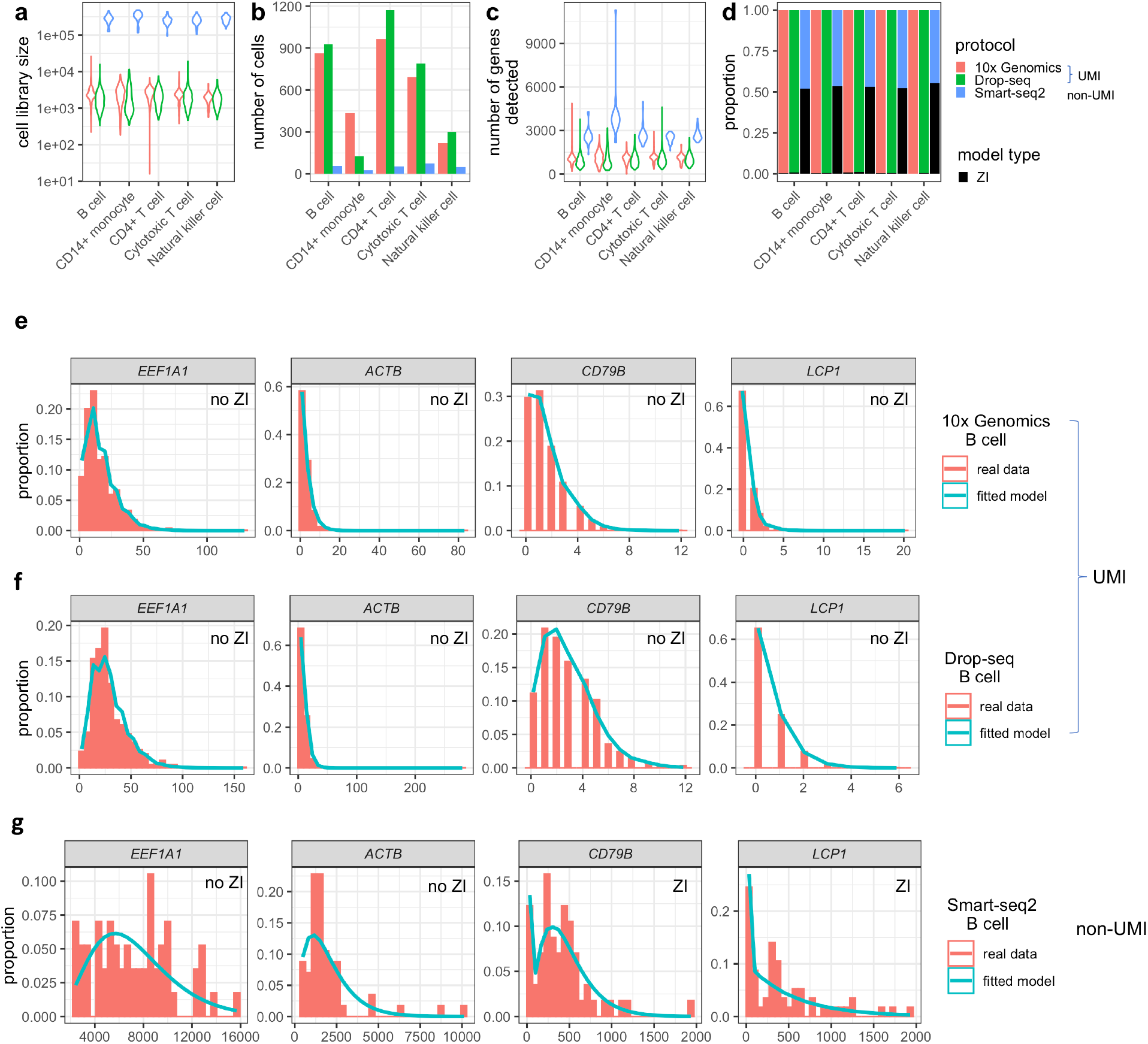
Statistical modeling of 10x Genomics, Drop-seq, and Smart-seq2 data for the same PMBC sample. 10x Genomics and Drop-seq data are UMI-based, while Smart-seq2 data are non-UMI-based. **(a)** Violin plots showing the distribution of cell library sizes for each of five PMBC cell types measured by each protocol [43]. For the UMI-based protocols (10x Genomics and Drop-seq) and Smart-seq2, a cell’s library size is defined as the total number of UMIs and reads, respectively, in that cell. **(b)** Barplots showing the number of cells detected for each cell type by each protocol. **(c)** Violin plots showing the distribution of the number of genes detected per cell. **(d)** Barplots showing the proportions of genes for which zero-inflated (ZI) models are chosen (black) and genes for which non-zero-inflated models are chosen (non-black). The model selection is done by likelihood ratio tests. **(e)** Four example genes’ distributions of UMI counts in B cells measured by 10x Genomics. The observed count distributions are shown in histograms. Non-zero-inflated (no ZI) models are chosen, and the fitted model distributions are shown in cyan curves. **(f)** The same four example genes’ distributions of UMI counts in B cells measured by Drop-seq. **(g)** The same four example genes’ distributions of read counts in B cells measured by Smart-seq2. ZI models are chosen for two genes.

Next, for each gene in each dataset, we fit the four distributions to its read or UMI counts in cells of each type, and we choose its distribution among the four distributions by likelihood ratio tests (see [102] for detail). The rationale is to choose the least complex distribution that fits the data well. Fig. 5d shows that non-zero-inflated distributions (Poisson and NB) are chosen for almost all genes in the 10x Genomics and Drop-seq datasets, while zero-inflated distributions (ZIP and ZINB) are chosen for about half of the genes in the Smart-seq2 dataset. This result is consistent with the recent advocate for not using zero-inflated models for UMI-based data [64, 94], and it suggests that zero-inflated modeling is still useful for Smart-seq2 data. For illustration purposes, in Fig. 5e–g, we plot the read or UMI count distributions for four genes (*EEF1A1, ACTB, CD79B*, and *LCP1*) in B cells in these three datasets, and we also plot the fitted chosen distribution for each gene. Specifically, non-zero-inflated distributions are chosen for all the four genes in the UMI-based datasets, while zero-inflated distributions are chosen for *CD79B* and *LCP1* in the Smart-seq2 dataset. Our results show that the same gene’s expression distribution under the same biological condition may be described by different statistical models for data generated by different protocols, confirming that zero inflation provides no direct information on biological zeros, whose existence does not depend on protocols (Fig. 1b).

### How non-biological zeros affect scRNA-seq data analysis

To evaluate the effects of non-biological zeros on scRNA-seq data analysis, such as cell clustering and DE gene identification, we need access to true cell types and true DE genes. Hence, we use scDesign2 [102], a probabilistic, flexible simulator we developed to generate realistic scRNA-seq count data from any protocol with gene correlations captured. First, we train scDesign2 on the three benchmark PBMC datasets (10x Genomics, Drop-seq, and Smart-seq2) [43], which all contain the same five cell types (B cells, CD14+ monocytes, CD4+ T cells, cytotoxic T cells, and natural killer cells) and are used in Fig. 5. Second, we simulate the corresponding non-zero-inflated synthetic datasets, one for each protocol, in the form of gene-by-cell count matrices. In detail, after the first training step, every gene in each cell type has a fitted count distribution (Poisson, ZIP, NB, or ZINB) by scDesign2; in the second simulation step, we generate read or UMI counts for every gene in each cell type from the non-zero-inflation component (Poisson or NB). Note that we set the number of synthetic cells generated by scDesign2 equal to the number of real cells for each cell type. Hence, for each gene, this simulation procedure removes the statistical zero inflation, which we define in the last section, and provides the gene’s expected expression level in each cell type (as the mean of its non-zero-inflation component).

The reason why we use scDesign2 [102] as the simulator is that we desire synthetic cells that preserve real genes and gene-gene correlations observed in real data. The reason is that real genes are the targets of DE analysis, and synthetic cells with realistic gene-gene correlations are necessary for evaluating cell clustering and dimension reduction. As discussed in the scDesign2 paper [102], simulators such as SymSim [103] and Splatter [68] do not preserve real genes, and Sergio [104] requires an additional input of a gene regulatory network and cannot preserve the observed gene-gene correlations in real data. We note that we do not aim to benchmark simulators in this work, so we choose scDesign2, which preserves genes and gene-gene correlations and allows us to generate non-zero-inflated data, making it easy for us to introduce non-biological zeros using various masking schemes.

To simulate scRNA-seq data with known DE genes, we first use scDesign2 to fit probabilistic models—selected from Poisson, NB, ZIP, and ZINB models—to each cell type in each real PBMC dataset generated by Smart-seq2, Drop-seq, or 10x Genomics. Then we focus on two cell types, CD4+ T cells and Cytotoxic T cells, and define the true DE genes as the top 1500 genes that have the largest estimated mean differences (in the fitted probabilistic models) between the two cell types. We consider the remaining genes as true non-DE genes, and we set the mean parameter of each true non-DE gene to be the same for the two cell types (by averaging the gene’s estimated mean parameters in its two fitted models for the two cell types). Finally, we use scDesign2 to generate synthetic scRNA-seq data without zero inflation: for every true DE gene, we generate its synthetic counts from its two fitted models (with zero inflated components removed if existent) for the two cell types; for every true non-DE gene, we generate its synthetic counts from two altered models (with the same averaged mean, zero inflation components removed if existent, and NB’s dispersion parameters kept as they are estimated in the two fitted models).

Based on the three non-zero-inflated synthetic datasets (10x Genomics, Drop-seq, and Smart-seq2), we define the positive controls for two typical analyses: cell clustering and DE gene identification, which are ubiquitous in scRNA-seq data analysis pipelines. For cell clustering, the positive controls are provided by scDesign2 as the cell types from which it generates synthetic cells. For DE gene identification, the positive controls are provided by scDesign2 as the genes whose expected expression levels differ between cell types.

Using each of the five masking schemes (see Additional file 1), we introduce a varying number of non-biological zeros, corresponding to masking proportions *p* = 0.1, …, 0.9, into the three synthetic datasets corresponding to 10x Genomics, Drop-seq, and Smart-seq2 protocols, creating three suites of zero-inflated datasets, one suite per protocol. Note that each suite contains one non-zero-inflated dataset and 45 = 9 (# of masking proportions) *×* 5 (# of masking schemes) zero-inflated datasets. Then we apply Monocle3 (R package version `0.2.3.0`) [28] and Seurat (R package version `3.2.1`) [13], two popular multi-functional software packages, to the three suites of datasets. We use the two packages to perform cell clustering and DE gene identification, and we evaluate the analysis results based on our previously defined positive controls. Fig. 6a–c summarizes how the accuracy of the two analyses deteriorates as the masking proportion increases under each masking scheme and for each protocol.

**Figure 6:**
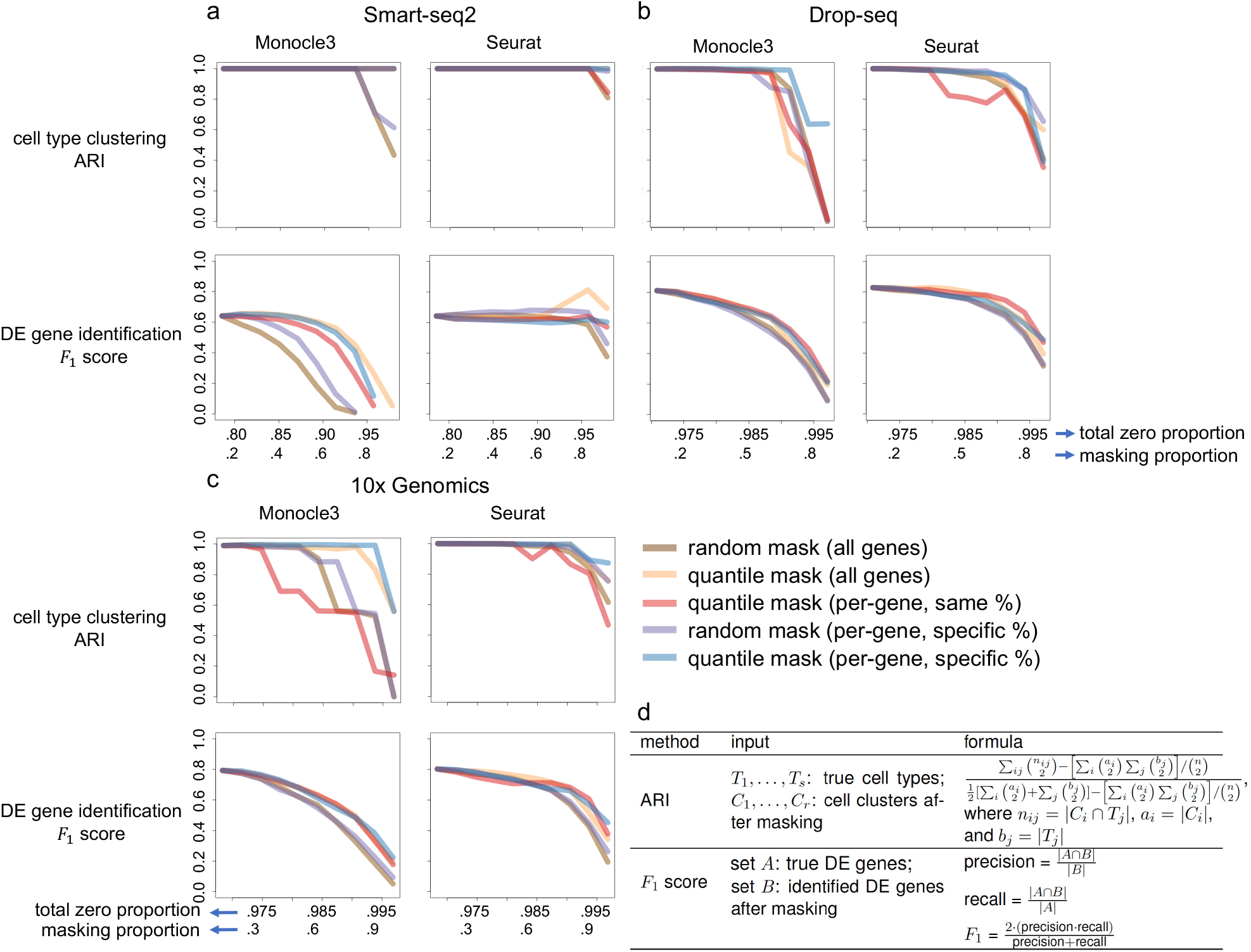
Effects of non-biological zeros on cell clustering and DE gene identification. We introduce a varying number of non-biological zeros, which correspond to masking proportions 0.1–0.9, into the simulated **(a)** Smart-seq2, **(b)** Drop-seq, and **(c)** 10x Genomics datasets using five masking schemes. The horizontal axes show (top) the total zero proportion (including the zeros before masking and the non-biological zeros introduced by masking) and (bottom) the masking proportion (i.e., the proportion of non-zero counts masked by a masking scheme). After introducing non-biological zeros, we apply Monocle 3 and Seurat to each dataset to perform cell clustering and identify DE genes. For the two analyses, we evaluate their accuracy using the adjusted rand index (ARI) and *F*_1_ score (given the false discovery rate 5%), respectively. **(d)** Technical definitions of the ARI and *F*_1_ score.

The clustering results (top rows in Fig. 6a–c) show that the clustering accuracy (measured by the adjusted rand index; Additional file 1: Fig. S1d) is robust to the introduction of non-biological zeros up to the masking proportion *p* = 0.6 (i.e., 60% non-zero counts are masked as zeros) for most masking schemes. Compared with Monocle3, Seurat is more robust to non-biological zeros under all the five masking schemes. Among all schemes, the two schemes that assume (1) dependence between masking and count values and (2) gene-specific masking proportions—`quantile mask (all genes)` and `quantile mask (per-gene, specific %)` (Additional file 1: Fig. S1b)—have the least deteriorating effects on cell clustering. This result is reasonable as these two schemes tend to mask low counts to zeros so that the relative order of gene expression counts (from low to high) is better preserved than by the other three schemes. A recent article argues that zeros in scRNA-seq data carry biological meanings and should be embraced, and its argument is based on the assumption that most zeros correspond to low expression levels [66], an assumption aligned with these two masking schemes. Finally, among the three protocols, clustering on Smart-seq2 data is most robust to non-biological zeros, likely because Smart-seq2 data contain fewer zeros than the two UMI-based protocols’ data do. It is worth noting that, between the two UMI-based protocols, clustering accuracy is better on 10x Genomics data than Drop-seq data.

The DE gene identification results (bottom rows in Fig. 6a–c) show that the *F*_1_ scores (at 5% false discovery rate; Fig. 6d) are robust to non-biological zeros for Seurat, but not as much for Monocle3. The reason is that Seurat uses MAST [36], a method built upon a zero-inflated model, for DE gene identification, while Monocle3 uses non-zero-inflated models (including Poisson, quasi Poisson, and NB) that cannot account for additional non-biological zeros. Among the five masking schemes, the two random schemes that assume independence between masking and count values—`random mask (all genes)` and `random mask (per-gene, specific %)` (Ad-ditional file 1: Fig. S1b)—have the most deteriorating effects on DE gene identification. This result is reasonable as these two schemes may mask high counts to zeros, so they disrupt every gene’s count distribution more than the other three schemes do. Interestingly, although `quantile mask (per-gene, same %)` is unlikely a realistic generation mechanism of non-biological zeros as it masks the same proportion of non-zero counts for every gene, we observe that Seurat has robust *F*_1_ scores as non-biological zeros are introduced by this scheme. This seemingly unexpected result reflects that zero-inflated models are robust for DE gene identification under quantile masking, even though the masking proportion may not be reasonable. Finally, regarding the three protocols, Seurat has better *F*_1_ scores for Smart-seq2 data than Monocle3 does, a reasonable result given the observed zero-inflation in Smart-seq2 data (Fig. 5d). For the two UMI-based protocols, Monocle3 and Seurat have comparable performance in terms of *F*_1_ scores. We have also observed that the DE analysis results for UMI-based data are better than for non-UMI-based data. One possible reason is the larger sample sizes (larger numbers of cells) in Drop-seq and 10x data that increase the power in statistical testing. To supplement the *F*_1_ scores, we show the corresponding precision and recall rates in Additional file 1: Fig. S2. It is worth noting that although the false discovery rate is set to 5%, the precision rates of both Monocle3 and Seurat are far below the expected precision 95%, which is equal to one minus the false discovery rate. This phenomenon calls for better false discovery rate control in scRNA-seq DE analysis [105]. In addition, compared to Seurat, Monocle3 shows a greater fluctuation in both precision and recall as the masking proportion increases.

In summary, compared with DE gene identification, cell clustering is more robust to non-biological zeros. This result suggests that the sparsity in scRNA-seq data affects gene-level analyses more than cell-level analyses because the latter jointly uses all genes’ expression levels. Overall, Seurat is more robust than Monocle3 is to non-biological zeros for both analyses. For cell clustering, Seurat has better accuracy regardless of protocols. For DE gene identification, Seurat is preferable for Smart-seq2 data, while Monocle3 has better accuracy for UMI-based data.

It is worth noting that many imputation methods evaluate their imputation accuracy based on only the `random mask (all genes)` scheme [49, 62, 106]. Our results indicate that non-biological zeros introduced by different masking schemes have different effects on cell clustering and DE gene identification, and quantile masking may be more realistic given previous reports that genes with lower expression values have more zeros than genes with higher expression [39, 46]. Hence, we urge that quantile masking schemes be considered in the future evaluation of computational methods that deal with non-biological zeros.

### Input data: observed vs. imputed vs. binarized counts

Current scRNA-seq data analysis typically takes three types of input data: observed, imputed, and binarized counts. Researchers use imputed and binarized counts to deal with the vast proportion of zeros. Although log-transformed counts are often used as input data, this practice is under controversy [93, 107, 108] and is not the focus of our discussion. Here we summarize the advantages, disadvantages, and suitable users (bioinformatics tool developers vs. users) of each input data type.

Direct modeling of observed counts is the most common practice for bioinformatics tool developers [13, 28, 30–33, 36, 109]. An obvious advantage of direct modeling is that observed counts are not biased by any data pre-processing steps. Hence, observed counts are the preferred input data type for most tool developers. However, unlike tool developers, tool users need to apply existing bioinformatics tools to scRNA-seq data. If the observed counts do not work well with existing tools, for practical reasons, tool users may consider data pre-processing steps such as imputation and binarization so that existing tools can output reasonable analysis results.

Since the sparsity in scRNA-seq counts has posed a great hurdle for many existing tools, imputation has been proposed as a practical data pre-processing step, and many imputation methods have been developed [20, 44, 45, 47–63]. Of course, imputation has the risk of biasing data, leading to false signals [99] or diminished biological variation [45, 63, 110]. For example, Chen *et al*. pointed out that a major drawback of scRNA-seq imputation is diminished gene expression variability across cells after imputation; they argued that it would be important to quantify the expression variability before and after imputation, in addition to evaluating the mean expression prediction (by, for example, comparing it to the gene expression measurement in the same cell type in a separate bulk dataset) [63]. However, imputation has two practical advantages for single-cell biologists who are mostly tool users. First, many imputation methods have shown that their imputed counts, in which many zeros in the observed counts become non-zeros, agree better with biological knowledge and/or biologists’ expectations. For example, the effectiveness of imputation has been supported by evidence that scRNA-seq data after imputation agree better with bulk RNA-seq data or single-cell RNA fluorescence in situ hybridization (FISH) data [48, 62, 111]. Second, imputation builds a bridge that connects sparse scRNA-seq data to many powerful tools designed for non-sparse data. For example, DESeq2 [35] and edgeR [31] are two popular DE gene identification methods for bulk RNA-seq data; however, they are not directly applicable to scRNA-seq data because their models do not account for data sparsity. Hence, if tool users cannot find a DE gene identification method that works well for their scRNA-seq data, they may consider reducing zeros by imputation methods to make DESeq2 or edgeR applicable [60, 61, 63], conditional on verified false discovery rate control [105, 112].

Moreover, a recent article provides a new perspective by proposing to use only binarized counts (with all non-zero counts truncated as ones) for cell clustering [66]. It argues that, by removing the magnitudes of non-zero counts, binarization alleviates the need for normalizing individual cells’ sequencing depths. Further, its key message is that zeros are biologically meaningful because binarized counts can lead to reasonable cell clustering results. Other works also suggest that binarized counts can serve as useful data, in addition to observed counts, and be incorporated into scRNA-seq data modeling and analysis [23, 113]. Although binarized counts eliminate the expression differences between highly- and lowly-expressed genes, they highlight the co-expression patterns of genes, i.e., whether two genes are co-expressed in a cell, which have been used in marker gene selection [12] and gene network construction [114–116]. However, it remains unclear whether binarized counts can replace observed counts in scRNA-seq data analysis. Our intuition says that the answer is unlikely yes for all analyses because the magnitudes of non-zero counts reflect expression levels of genes in each cell. Qiu uses binarized counts to deal with cell clustering, a cell-level analysis [66]. For gene-level analyses such as DE gene identification, binarized counts are unlikely better than observed counts. For example, if a gene has similar percentages of zero counts in two cell types, but its non-zero counts are much larger in one cell type than the other, then this gene should be identified as DE using observed counts, but it would be missed as DE using binarized counts. In the previous section “How non-biological zeros affect scRNA-seq data analysis,” we have compared the effects of non-biological zeros on clustering and DE gene analysis. For tool developers, it would be beneficial to consider using binarized counts in addition to observed counts for developing new analysis tools. For tool users, binarized counts can be used for exploratory data analysis because several efficient computational tools are applicable to binary counts only, e.g., scalable probabilistic principal component analysis [117].

We further evaluate the effects of the three input data types (observed, binarized, and imputed counts) on three popular downstream analyses: cell clustering, cell dimension reduction (two-dimensional visualization), and DE gene identification. To obtain the imputed counts, we use three popular imputation methods: scImpute, MAGIC, and SAVER, which demonstrate good performance in a recent study that benchmarked 18 imputation methods [118]. We note that our goal here is not to benchmark the existing *>* 70 imputation methods (https://www.scrna-tools.org/tools?sort=name&cats=Imputation) but to demonstrate the importance of considering various zero generation processes (i.e., masking schemes) to achieve fair benchmarking of computational methods. For readers interested in evaluating other imputation methods, we have released our code for the five masking schemes (Additional file 1) on GitHub (https://github.com/ruochenj/Five_masking_schemes).

To benchmark cell clustering and dimension reduction results, we use the three real scRNA-seq PBMC datasets with labelled cell types [43]—one non-UMI-based dataset generated by Smart-seq2 and two UMI-based datasets generated by 10x Genomics and Drop-seq—which we have used in the previous section to evaluate the effects of non-biological zeros. To benchmark DE gene identification results, we generate synthetic datasets containing pre-defined true DE genes by scDesign2 [102] from two cell types (CD4+ T cells and cytotoxic T cells) in the three real datasets. (The definition of DE genes is described in the previous section.)

For cell clustering, we use two algorithms: Qiu’s algorithm designed specifically for binarized counts [66] and the Louvain algorithm (implemented in Seurat). For all three input data types, we use the Louvain algorithm to cluster cells; for binarized counts only, we also use Qiu’s algorithm. Based on the cell type labels provided in all three datasets, we calculate the ARI as a measure of clustering accuracy (Additional file 1: Fig. S3; top row).

For cell dimension reduction (two-dimensional visualization), we perform PCA, t-SNE and UMAP (implemented in Seurat) on the observed and imputed counts, and then we use the average Silhouette score to evaluate how well the labeled cell types are separated in the two-dimensional space (Additional file 1: Fig. S4). We observe that UMAP has the overall best performance and thus decide to use UMAP to evaluate the impacts of imputation methods and binarization on the dimension reduction analysis (Additional file 1: Fig. S5). Additional file 1: Figs. S6, S7, and S8 show UMAP’s two-dimensional reduction results of Smart-seq2, Drop-seq, and 10x Genomics data, respectively.

For DE gene analysis, we consider two DE methods. For all three inpute data types, we use MAST (implemented in Seurat) to perform DE gene identification; for binarized counts only, we also apply a two-sample proportion test to the binarized data. At a 5% false discovery rate, we use precision (Additional file 1: Fig. S9), recall (Additional file 1: Fig. S10) and F_1_ score (Additional file 1: Fig. S11) to evaluate the identification results.

Fig. 7a–b summarizes the relative performance of the three input data types for the three protocols (Smart-seq2, Drop-seq, and 10x Genomics) in the three downstream analyses. In terms of cell clustering, for non-UMI-based Smart-seq2 data, the Louvain algorithm has better performance on scImpute and SAVER’s imputed counts than on the observed or binarized counts; for UMI-based Drop-seq and 10x Genomics data, the Louvain algorithm on the observed counts and Qiu’s algorithm on the binarized counts have comparable performance and outperform the Louvain algorithm applied to other input data types, suggesting that imputation does not improve the clustering of UMI-based data. Notably, Qiu’s algorithm only works well for binarized counts of UMI-based data, likely due to its special design. In terms of cell dimension reduction, scImpute’s imputed counts work the best for non-UMI-based data; the observed counts have the best performance for UMI-based data; binarized counts and MAGIC’s imputed counts have poor performance for both non-UMI-based and UMI-based data. In terms of DE gene analysis, for non-UMI-based data, all three imputation methods’ imputed counts outperform the observed and binarized counts, a result consistent with our previous discussion on the existence of zero-inflation in non-UMI-based data; for UMI-based data, the observed counts and scImpute’s imputed counts lead to the best result for Drop-seq and 10x Genomics data, respectively.

**Figure 7:**
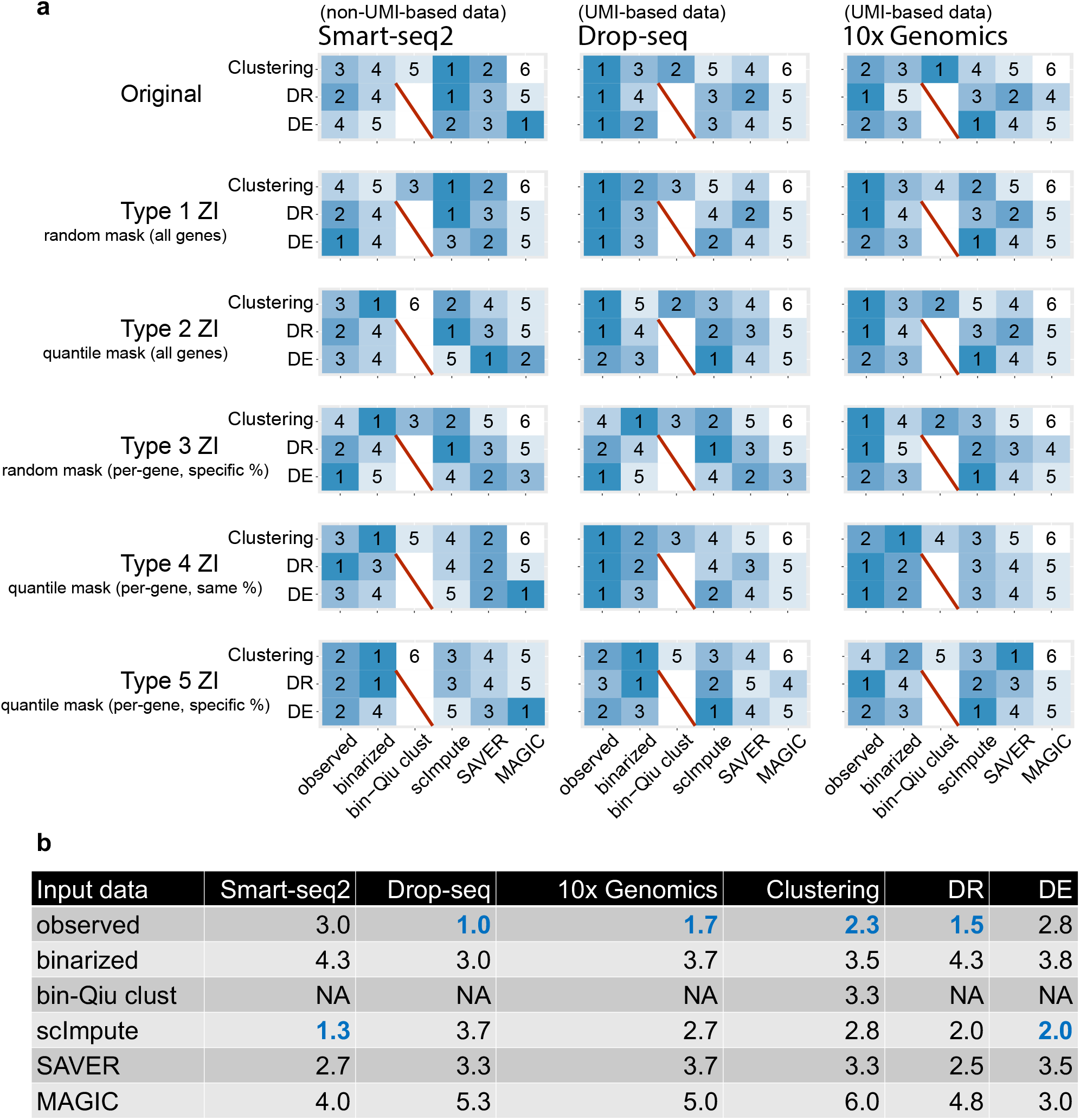
Performance ranks of using observed, binarized, and imputed counts (from three experimental protocols ad under five masking schemes) in three downstream analyses. **(a)** We perform cell clustering (Clustering), cell dimension reduction (DR), and gene differential expression (DE) analysis on the observed, binarized, and imputed counts of Smart-seq2, Drop-seq, and 10x Genomics data. We consider three popular imputation methods: scImpute, SAVER, and MAGIC. In addition to the original data, we use five masking schemes (Type 1 ZI–Type 5 ZI) to introduce 50% non-biological zeros and evaluate the effects on the downstream analyses with different input data. The five masking schemes are `random mask (all genes)`, `quantile mask (all genes)`, `random mask (per-gene, specific %)`, `quantile mask (per-gene, same %)`, and `quantile mask (per-gene, specific %)`, corresponding to type 1 ZI–type 5 ZI, respectively. The six columns correspond to different input data types: observed counts, binarized counts, binarized counts analyzed by the Qiu’s clustering algorithm (bin-Qiu clust), imputed counts by scImpute, imputed counts by SAVER, and imputed counts by MAGIC. For cell clustering, except bin-Qiu clust, clustering is conducted by the Louvain clustering algorithm (in Seurat); clustering performance is ranked by the ARI. For cell DR analysis, we apply UMAP (in Seurat) to perform DR and calculate the average Silhouette score (based on known cell types) for each input data type to evaluate the DR performance. For gene DE analysis, we apply the two-sample proportion test to binarized counts and MAST (in Seurat) to observed, binarized, and imputed data to perform DE analysis. To rank the DE performance by the *F*_1_ score (at the 5% false discovery rate), since binarized counts have two DE methods, we compute the rank for the better-performing method in each comparison. In each row of each matrix, rank 1 indicates the best-performing input data type, while rank 6 indicates the worst. **(b)** On the original data, we compute the average ranks of the six input data types. Columns 4–6 show the average ranks for Smart-seq2 data, Drop-seq data, and 10x Genomics data across the three downstream analysis—cell clustering analysis, cell DR analysis, and gene DE analysis. Columns 7–9 show the weighted averages of the ranks for the three downstream analysis given the three protocols. Weights of 2, 1, 1 are used for Smart-seq2, Drop-seq, and 10x Genomics to ensure that the weights for non-UMI-based and UMI-based data are equal.

Moreover, we evaluate the three input data types in the three downstream analyses after applying the five masking schemes (see Additional file 1) to introducing additional non-biological zeros. Under each masking scheme, we mask 50% of the original non-zero counts as zeros in each of the three original datasets (Smart-seq2, Drop-seq, and 10x Genomics). In terms of cell clustering analysis, for non-UMI-based data, scImpute’s imputed counts demonstrate robust performance and stay as a top-performing input data type under the first three masking schemes, including the two random masking schemes; interestingly, by the Louvain algorithm, the binarized counts do not perform well for the original data but become a top-performing input data type under the last four masking schemes, including the three quantile masking schemes. These two results suggest that scImpute’s imputation and binarization ameliorate the effects of additional non-biological zeros in complementary ways. For UMI-based data, the observed counts lead to the overall best clustering results under all masking schemes (ranked the 1st in 6 out of 10 protocol-masking scheme combinations), while Qiu’s algorithm is not robust to the introduction non-biological zeros by masking schemes. In terms of cell dimension reduction, scImpute’s imputed counts are the best input data type for non-UMI-based data (ranked the 1st under 3 out of 5 masking schemes), while the observed counts are the best for UMI-based data (ranked the 1st in 8 out of 10 protocol-masking scheme combinations). In terms of DE gene analysis, for non-UMI-based data, there is no universal winner: the observed counts work the best under random masking schemes, while SAVER and MAGIC’s imputed counts work the best under quantile masking schemes. For UMI-based data, scImpute’s imputed counts have the best performance (ranked the 1st in 6 out of 10 protocol-masking scheme combinations), followed by the observed counts (ranked the 1st in 4 out of 10 protocol-masking scheme combinations).

In summary, the observed counts work well for UMI-based data and are robust to the introduction of non-biological zeros. As expected, the binarized counts work well under the quantile masking schemes, which largely preserve the ranks of gene expression levels. Qiu’s clustering algorithm works well for the binarized counts of UMI-based data but is not robust to the introduction of non-biological zeros. Imputation methods show concrete improvement for non-UMI-based data, but not so much for UMI-based data. This is consistent with the findings by Kim *et al*. that imputing UMI-based data can introduce unwanted noise and is thus not recommended [65]. Among the imputation methods, scImpute shows the best performance, while MAGIC does not perform well; a likely reason is that the data we use contain discrete cell types instead of continuous cell trajectories. Notably, the performance of imputation methods depends heavily on the masking scheme, demonstrating the importance of considering multiple masking schemes for the development and benchmarking of imputation methods.

### Future directions

ScRNA-seq technologies have advanced the revelation of genome-wide gene expression profiles at the cell level. Accordingly, many computational algorithms and statistical models have been developed for analyzing scRNA-seq data. A well-known challenge in scRNA-seq data analysis is the prevalence of zeros, and how to best tackle zeros remains a controversial topic. Modeling and analysis may be performed on observed, imputed, or binarized scRNA-seq counts. However, the relative advantages and disadvantages of these three strategies remain ambiguous. In this article, we attempt to address this controversy by discussing multiple intertwined topics: the biological and non-biological sources of zeros, the relationship between zero prevalence and scRNA-seq technologies, the extent to which zero prevalence affects various analytical tasks, and the three strategies’ relative advantages, disadvantages, and suitable users. We benchmark the performance of analytical tasks on observed, binarized and imputed data with or without non-biological zeros introduced.

The prevalence of biological and non-biological zeros is a mixed result of intrinsic biological nature and complex scRNA-seq experiments. In particular, the generation mechanism of non-biological zeros is protocol dependent. Hence, it is infeasible to distinguish non-biological zeros from biological zeros purely based on observed counts. As a result, existing imputation methods have a glass ceiling if they use only observed counts as input. To better distinguish non-biological zeros from biological zeros, researchers need to utilize spike-in RNA molecules, whose true counts are known (e.g., External RNA Control Consortium spike-ins [119]), to investigate the generation mechanism of non-biological zeros. Such investigation requires consortium efforts such as the work by the Sequencing Quality Control (SEQC-2) consortium [120]. With a better understanding of how the generation of non-biological zeros depends on mRNA sequence features such as GC contents, statistical and mechanistic models may be developed to better distinguish non-biological zeros from biological zeros and thus to improve imputation accuracy.

The prevalence of biological and non-biological zeros is only one of the many obstacles in using scRNA-seq data for scientific discoveries. As scientific discovery is a trial-and-error process, scRNA-seq data analysis is unavoidably multi-step. Hence, bioinformatics tool developers must consider the pre-processing steps applied to input data and the downstream analyses users may perform on output data. Taking the popular Seurat package as an example, many data pre-processing steps are used before DE gene identification. These steps include filtering low-quality genes and cells, data normalization, gene selection, cell dimension reduction, and cell clustering. Hence, if tool developers are not aware of these pre-processing steps, their bioinformatics tools may not fit into the state-of-the-art scRNA-seq data analysis pipelines. Ultimately, the transparency and reproducibility of scRNA-seq data analysis call for a community collaboration between tool developers and users. Towards this goal, every research article, regardless of being tool development or data analysis, should contain a detailed description of each step and the underlying justifications [121].

## Availability of data and materials

The R code (license: Creative Commons Attribution 3.0 United States) for reproducing the results in Figs. 2–6 is available at https://doi.org/10.5281/zenodo.4393040 [122].

## Additional files

### Additional file 1

Supplementary material. It includes a detailed description of the five masking schemes, which introduce non-biological zeros to a scRNA-seq count matrix, and supplementary figures.

## Authors’ contributions

RJ and JJL designed the research. RJ conducted the research. RJ and TS wrote the computer code. JJL supervised the project. RJ, TS, DS, and JJL discussed the results and wrote the manuscript. All authors read and approved the final manuscript.

## Acknowledgements

The authors would like to thank the comments and feedback from Ms. Yiling Chen, Mr. Guanao Yan, Mr. Han Cui, Mr. Xinzhou Ge, and other members of the Junction of Statistics and Biology at UCLA (http://jsb.ucla.edu).

