## Additional file 1 for "Statistics or biology: the zero-inflation controversy about scRNA-seq data"

### Design of five masking schemes

Given the three synthetic datasets without zero inflation, we use five masking schemes to introduce a varying number of non-biological zeros into each dataset. Since there is no consensus on the generation mechanism of non-biological zeros, we design the five masking schemes to reflect two fundamental questions: whether the occurrence of non-biological zeros (1) depends on the actual gene expression levels and/or (2) is gene-specific. As the five masking schemes cover the extreme answers to both questions (Additional File 1: Fig. S1a), we expect that they together cover the unknown generation mechanism of non-biological zeros and would thus reveal the realistic effects of non-biological zeros on cell clustering and DE gene identification.

We provide a toy example to demonstrate the five masking schemes in Additional File 1: Fig. S1b and summarize their technical details in Additional File 1: Fig. S1c. In short, for a dataset with $n$ non-zero counts, given a masking proportion $p$, all schemes would mask approximately $np$ non-zero counts. However, the five schemes differ in masking which $np$ non-zero counts, and they can be categorized in two ways corresponding to the two aforementioned questions.

The first categorization is whether masking depends on the non-zero count values: random masking vs. quantile masking. While the two random masking schemes assume the independence between whether a non-zero count would be masked and the count value itself, the three quantile masking schemes assume a complete dependence by truncating non-zero values below a quantile (which corresponds to the masking proportion) to zero. Specifically, the two random masking schemes differ in the definition of independence: `random mask (all genes)` assumes the complete independence between masking and count values; `random mask (per-gene, specific %)` only assumes the conditional independence between masking and count values given each gene, and the masking proportion is gene-specific. Note that we define each gene's specific masking proportion as a function of the gene's non-zero counts based on an empirical formula

[1] Department of Statistics, University of California, Los Angeles, CA 90095-1554
[2] Bioinformatics Interdepartmental Ph.D. Program, University of California, Los Angeles, CA 90095-7246
[3] Department of Human Genetics, University of California, Los Angeles, CA 90095-7088
[4] Department of Computational Medicine, University of California, Los Angeles, CA 90095-1766

in the literature [1, 2] (Additional File 1: Fig. S1c); in short, the larger a gene's non-zero counts are, the smaller the gene's masking proportion is. Besides the two random masking schemes, the three quantile masking schemes differ in how they perform the truncation: `quantile mask (all genes)` truncates the lowest $100p\%$ non-zero counts of all genes; `quantile mask (per-gene, same %)` truncates the lowest $100p\%$ non-zero counts of each gene; `quantile mask (per-gene, specific %)` truncates the lowest non-zero counts of each gene based on the gene's specific masking proportion determined by the empirical formula.

The second categorization is regarding whether the masking proportion is gene-specific. Two schemes mask the same expected proportion $100p\%$ of non-zero counts for all genes: `random mask (all genes)` and `quantile mask (per-gene, same %)`. Three schemes use gene-specific masking proportions: `quantile mask (all genes)`, `random mask (per-gene, specific %)`, and `quantile mask (per-gene, specific %)`. Specifically, although `quantile mask (all genes)` does not use the empirical formula to determine gene-specific masking proportions as in `random mask (per-gene, specific %)` and `quantile mask (per-gene, specific %)`, it still truncates different proportions of non-zero counts for different genes. The reason is that its truncation threshold is set to the $p$-th quantile of all genes' non-zero counts, and different genes have different numbers of non-zero counts below that threshold. It is also worth noting that we do not include `random mask (per-gene, same %)` because it is theoretically equivalent to `random mask (all genes)`—both schemes are expected to randomly mask $100p\%$ of every gene's non-zero counts (Additional File 1: Fig. S1a).

Note that random masking aims to reflect the random nature of sampling zeros. In a sequencing experiment, allocation of reads to genes is essentially random sampling from a multinomial distribution, whose probabilities are the proportions of genes in terms of cDNA copy numbers in the sequencing library. Due to the randomness of sampling, for two genes with moderately different non-zero proportions, it is possible that, in one experiment, the gene with the larger proportion receives a zero read count, i.e., a sampling zero, while the gene with the smaller proportion receives a non-zero read count. The magnitude of the randomness depends on the sequencing depth. For every gene, the standard deviation of its count over its expected count is equal to a large constant depending on its proportion (i.e., $\sqrt{(1-q_i)/q_i}$, where $q_i$ is the proportion of gene $i$) multiplied by the inverse of the square root of the sequencing depth (i.e., $1/\sqrt{N}$, where $N$ is the sequencing depth). Hence, the smaller the sequencing depth, the larger the standard deviation of every gene's count in relation to its expected count, the more likely that genes receive sampling zeros irrespective of their proportions. Moreover, the expected number of sampling zeros (i.e., $\sum_{i=1}^{I}(1-q_i)^N$, where $I$ is the number of genes) decreases as the sequencing depth increases. In contrast, quantile masking aims to reflect gene proportions in the sequencing library and technical zeros, i.e., zero counts due to zero proportions without randomness. Quantile masking also reflects the fact that, despite of randomness, a gene with a small proportion is more likely to receive a sampling zero than a gene with a much larger proportion does.

Hence, for Drop-seq and 10x Genomics, since they sequence many cells, per-cell sequencing depth is low and thus randomness is influential, random masking better represents the occurrence of non-biological zeros, sampling zeros in particular, than quantile masking does. For Smart-seq2, since per-cell sequencing depth is high and thus randomness is negligible, quantile masking better resembles the generation mechanism of non-biological zeros, technical zeros in particular, than random masking does.

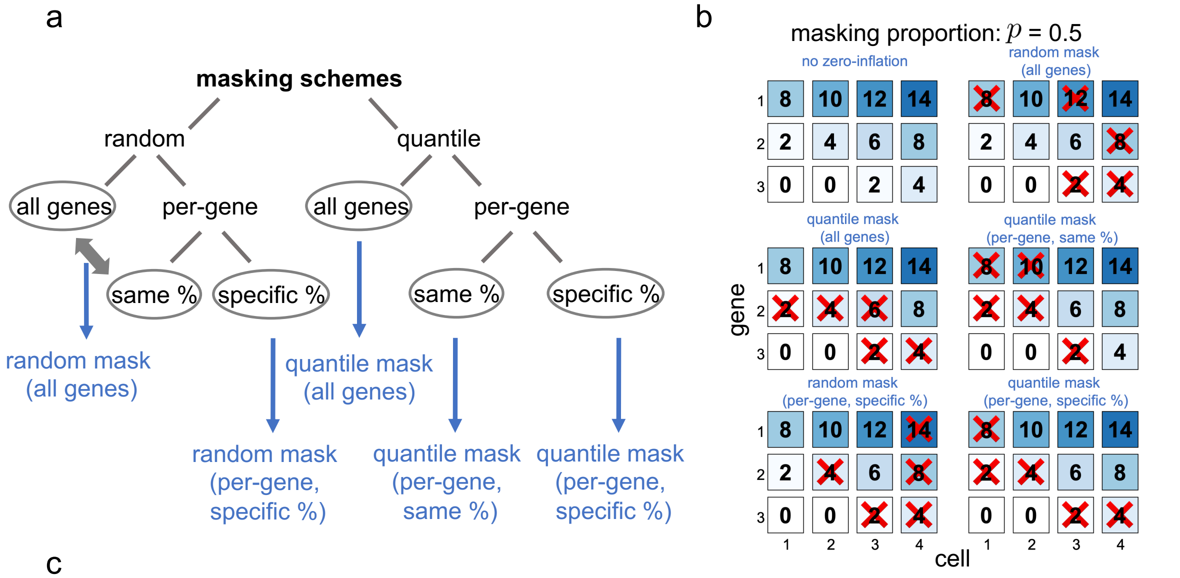


c

| masking scheme | parameter | method |
|---|---|---|
| random mask (all genes) | $p \in (0, 1)$ | randomly mask $N \sim$ Binomial$(n, p)$ of the $n$ non-zero counts of all genes |
| quantile mask (all genes) | $p \in (0, 1)$ | mask the smallest $\lfloor np \rfloor$ (floor) non-zero counts |
| qauntile mask (per-gene, same %) | $p \in (0, 1)$ | mask the smallest $\lfloor n_i p \rfloor$ non-zero counts of gene $i$, which has $n_i$ non-zero counts |
| random mask (per-gene, specific %) | $\lambda > 0, p \in (0, 1)$ | randomly mask $N_i \sim$ Binomial$(n_i, p_i)$ of the $n_i$ non-zero counts of gene $i$, whose average (natrual) log-transformed non-zero count is $\mu_i$, and $p_i = \exp(-\lambda \mu_i^2)$. $\lambda$ is chosen to satisfy $\sum_{i=1}^{\text{\# of genes}} n_i p_i = np$, where $n$ is the total number of non-zero counts of all genes |
| quantile mask (per-gene, specific %) | $\lambda > 0, p \in (0, 1)$ | mask the smallest $\lfloor n_i p_i \rfloor$ non-zero counts of gene $i$. $n_i$ and $p_i$ are the same as in "random mask (per-gene, specific %)" |

Figure S1: **Five masking schemes for introducing non-biological zeros.** **(a)** A tree diagram illustrating the design of the five masking schemes. From the top, the first division is about whether masking is independent of or completely dependent on count values, with the former as random masking and the latter as quantile masking. The second division is about whether masking is performed across all genes (with the same masking proportion) or within each gene (i.e., per-gene). If the latter, the third division is regarding whether the masking proportion is the same for all genes or specific to each gene depending on the gene's mean non-zero expression level. Note that random masking across all genes is equivalent to random masking per-gene with the same masking proportion (shown by the double arrow on the left). **(b)** A toy example illustration of the five masking schemes. The topleft plot shows the expression counts of three genes in four cells without zero-inflation; the other five plots show the expression counts after the five masking schemes are applied with the same masking proportion $p = 0.5$ (i.e., $50\%$ of the non-zero gene expression counts are masked as zeros). **(c)** Technical explanation of each masking scheme. In the notations, $p$ denotes the overall masking proportion across all genes, and $p_i$ is the masking proportion of gene $i$.

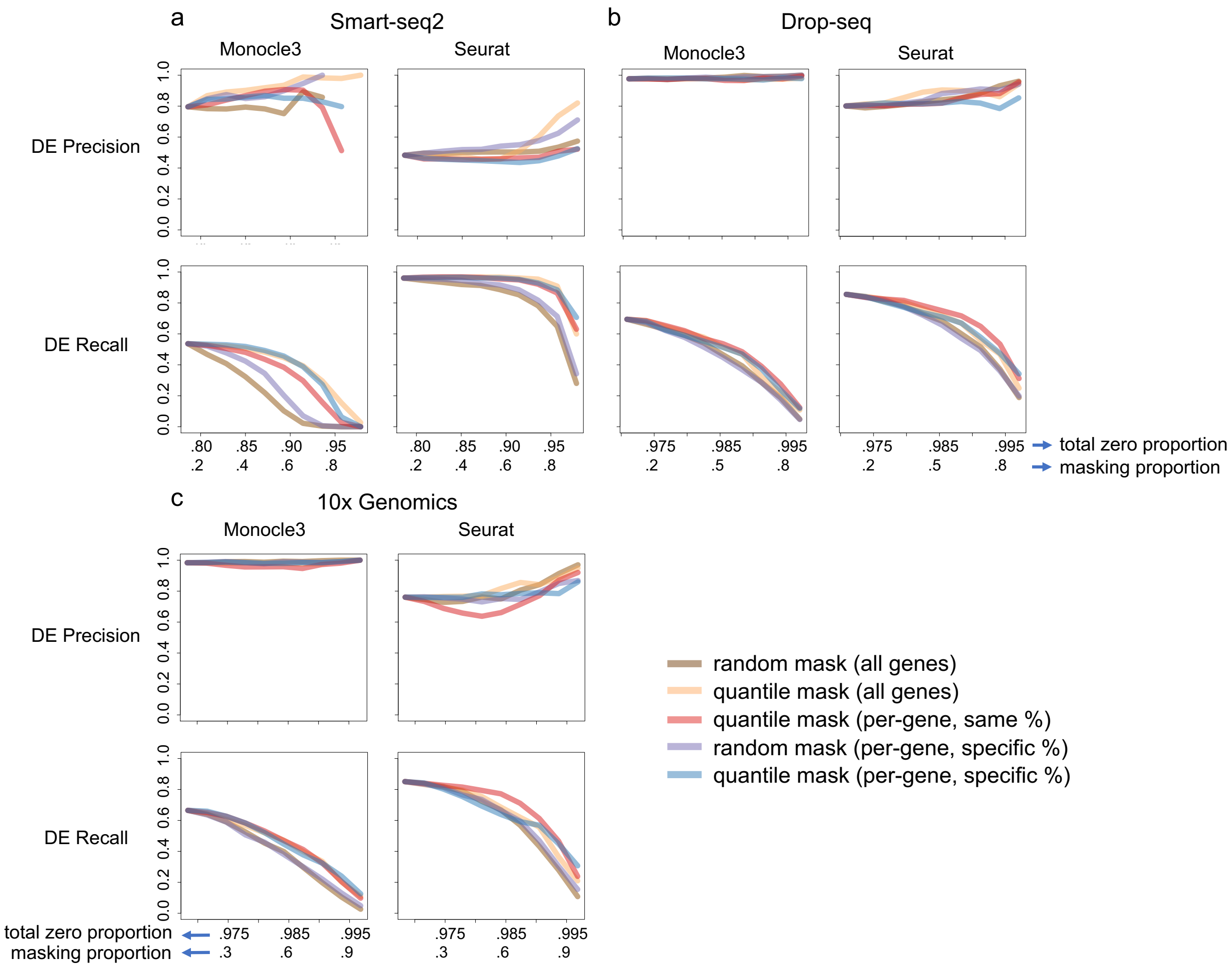


Figure S2: **Effects of non-biological zeros on DE gene identification in terms of precision and recall.** We introduce a varying number of non-biological zeros, which correspond to masking proportions $0.1$–$0.9$, into the simulated **(a)** Smart-seq2, **(b)** Drop-seq, and **(c)** 10x Genomics datasets using five masking schemes. The horizontal axes show (top) the total zero proportion (including the zeros before masking and the non-biological zeros introduced by masking) and (bottom) the masking proportion (i.e., the proportion of non-zero counts masked by a masking schemes). After the introduction of non-biological zeros, we apply Monocle 3 and Seurat to each dataset to identify DE genes. We evaluate the accuracy using the precision and recall (given the false discovery rate $5\%$; defined in Fig. **??**d), respectively.

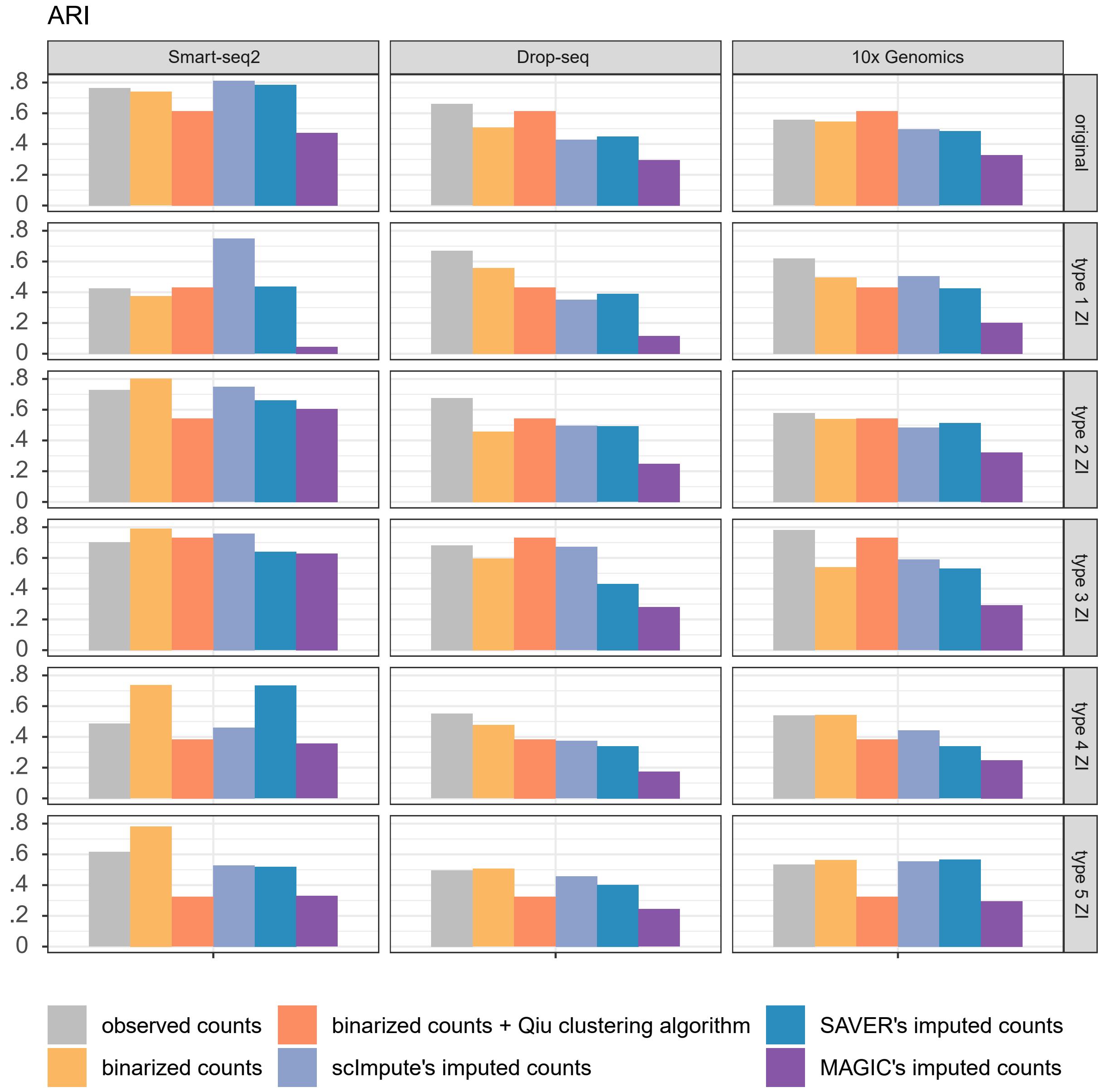


Figure S3: **Evaluation of clustering analysis on observed, binarized and imputed data.** We evaluate the clustering analysis on Smart-seq2, Drop-seq, and 10x Genomics data based on observed, binarized and imputed data. We perform this analysis before and after using the five masking schemes (type 1 ZI–type 5 ZI) to introduce non-biological zeros. Besides the bin-Qiu *et al.* which indicates the clustering algorithm developed specially for binarized data, we use Louvain clustering (in Seurat) on observed, binarized, and imputed data. We use ARI to evaluate the clustering results.

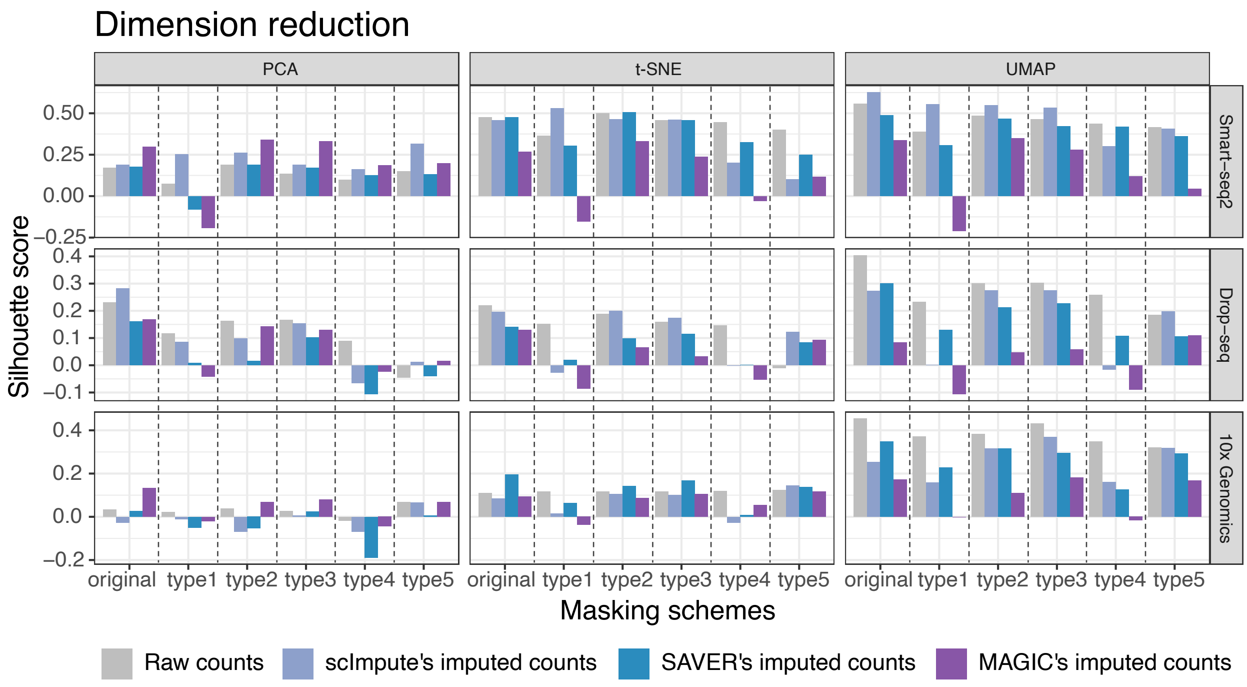


Figure S4: **Dimension reduction comparison using PCA, TSNE and UMAP.** In addition to the original data, we use five masking schemes (Type 1–Type 5) to introduce 50% non-biological zeros and evaluate the effects on the downstream analyses with different input data. The five masking schemes are `random mask (all genes)`, `quantile mask (all genes)`, `random mask (per-gene, specific %)`, `quantile mask (per-gene, same %)`, and `quantile mask (per-gene, specific %)`, corresponding to type 1–type 5, respectively. Then, we perform scImpute, Saver, or Magic to get the imputed data. Finally, We evaluate the dimension reduction using PCA, t-SNE or UMAP on Smart-seq2, Drop-seq, and 10x Genomics data based on observed and imputed data and use Silhouette score to evaluate the dimension reduction results.

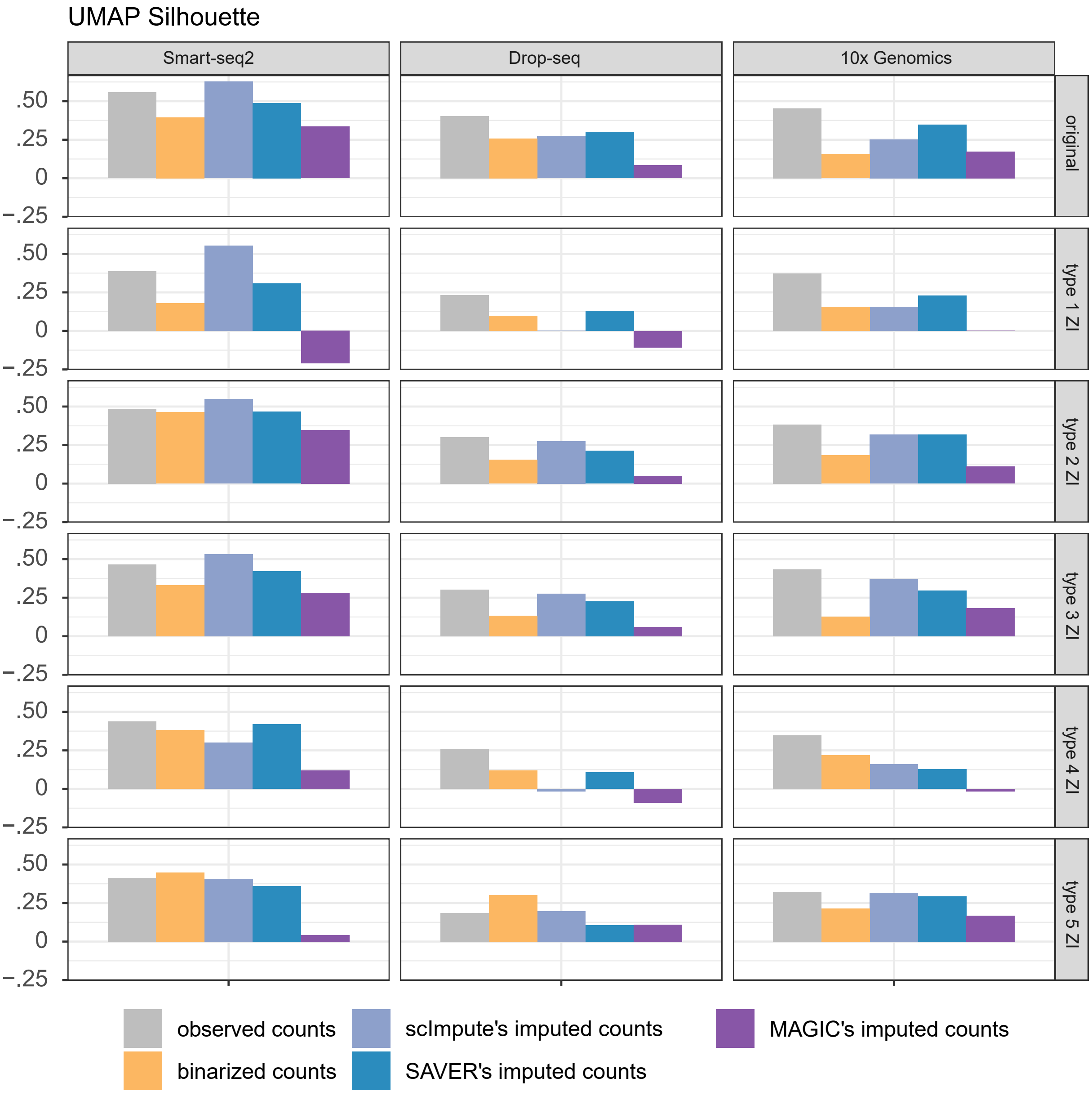


Figure S5: **Evaluation of dimension reduction analysis on observed, binarized and imputed data.** We evaluate the dimension reduction analysis on Smart-seq2, Drop-seq, and 10x Genomics data based on observed, binarized and imputed data. We perform this analysis before and after using the five masking schemes (type 1 ZI–type 5 ZI) to introduce non-biological zeros. We use UMAP (in Seurat) on observed, binarized, and imputed data to perform dimension reduction. We use Silhouette score to evaluate the dimension reduction results.

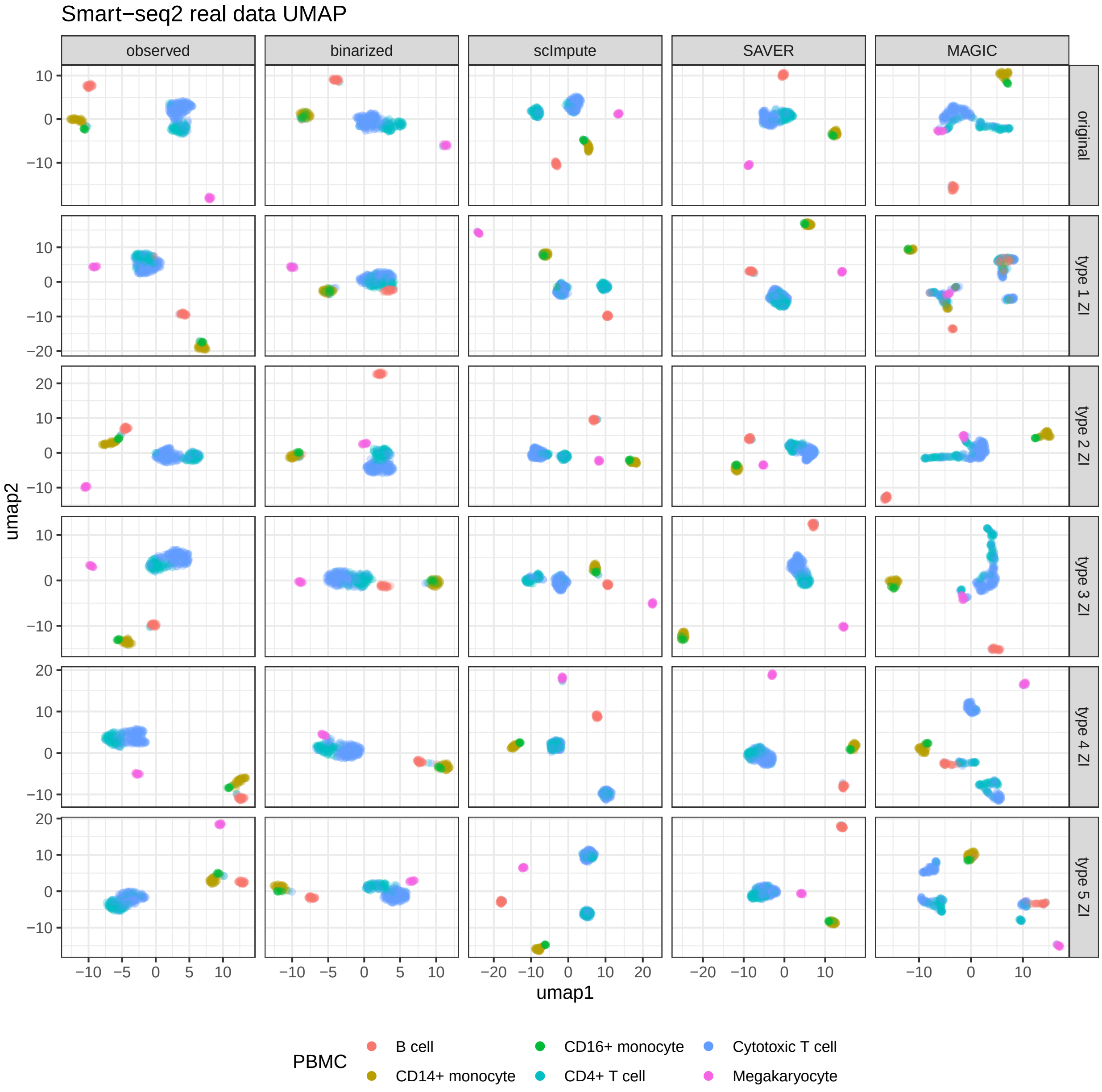


Figure S6: **UMAP dimesion reduction visualization on observed, binarized and imputed Smart-seq2 data.** We perform UMAP (in Seurat) on Smart-seq2's observed, binarized and imputed data. We perform this analysis before and after using the five masking schemes (type 1 ZI–type 5 ZI) to introduce non-biological zeros.

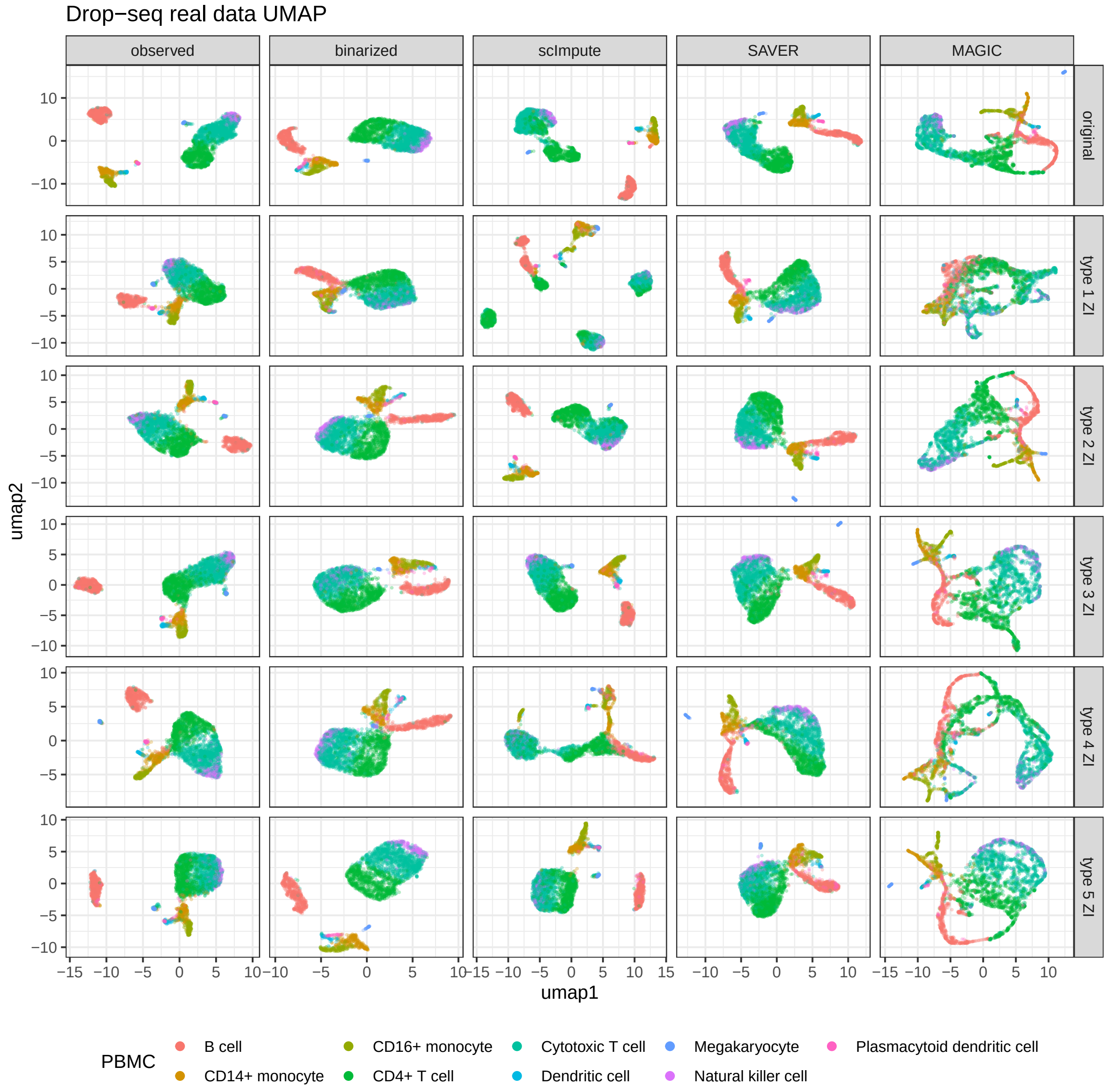


Figure S7: **UMAP dimesion reduction visualization on observed, binarized and imputed Drop-seq data.** We perform UMAP (in Seurat) on Drop-seq's observed, binarized and imputed data. We perform this analysis before and after using the five masking schemes (type 1 ZI–type 5 ZI) to introduce non-biological zeros.

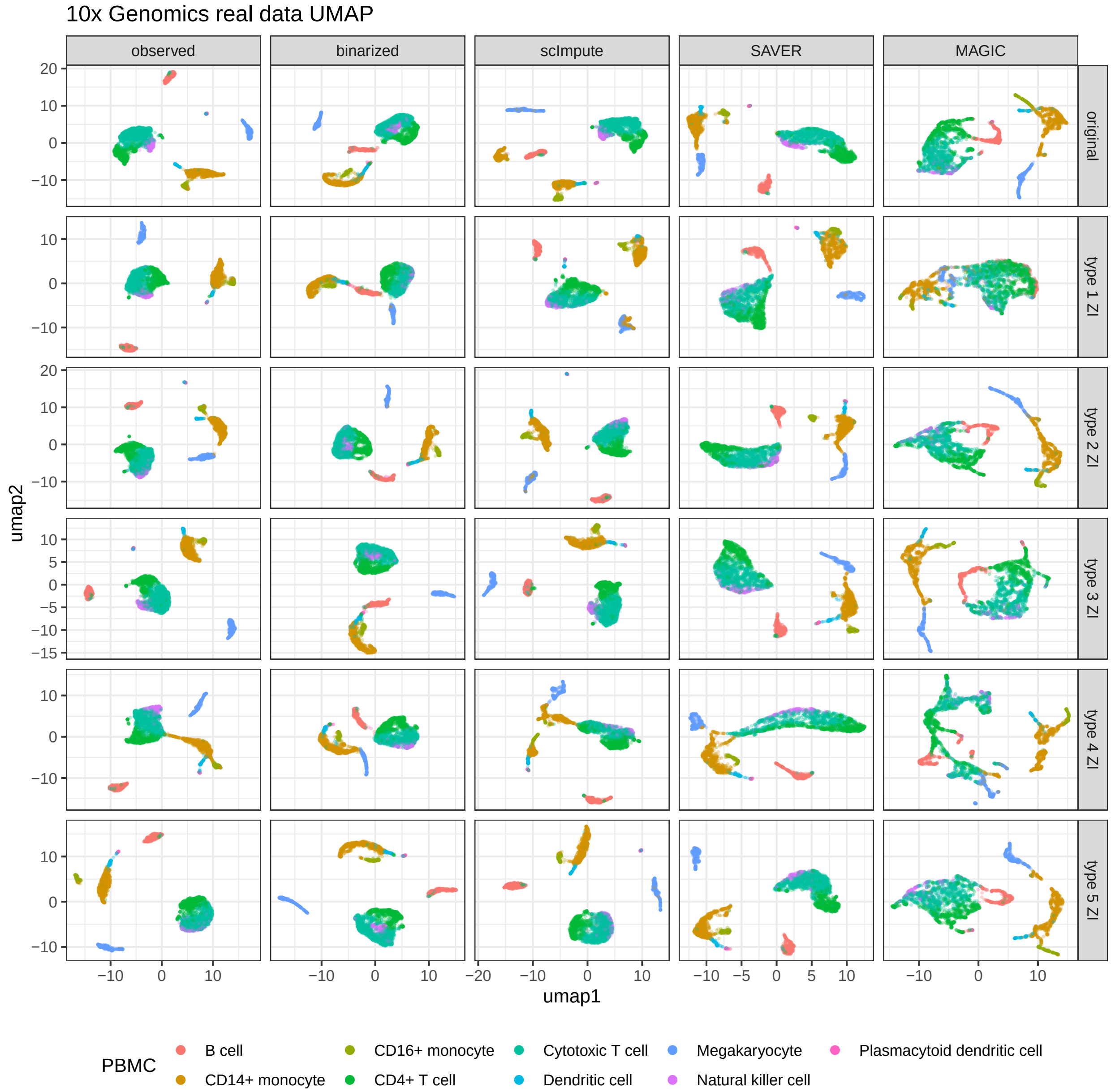


Figure S8: **UMAP dimesion reduction visualization on observed, binarized and imputed 10x Genomics data.** We perform UMAP (in Seurat) on 10x Genomics' observed, binarized and imputed data. We perform this analysis before and after using the five masking schemes (type 1 ZI–type 5 ZI) to introduce non-biological zeros.

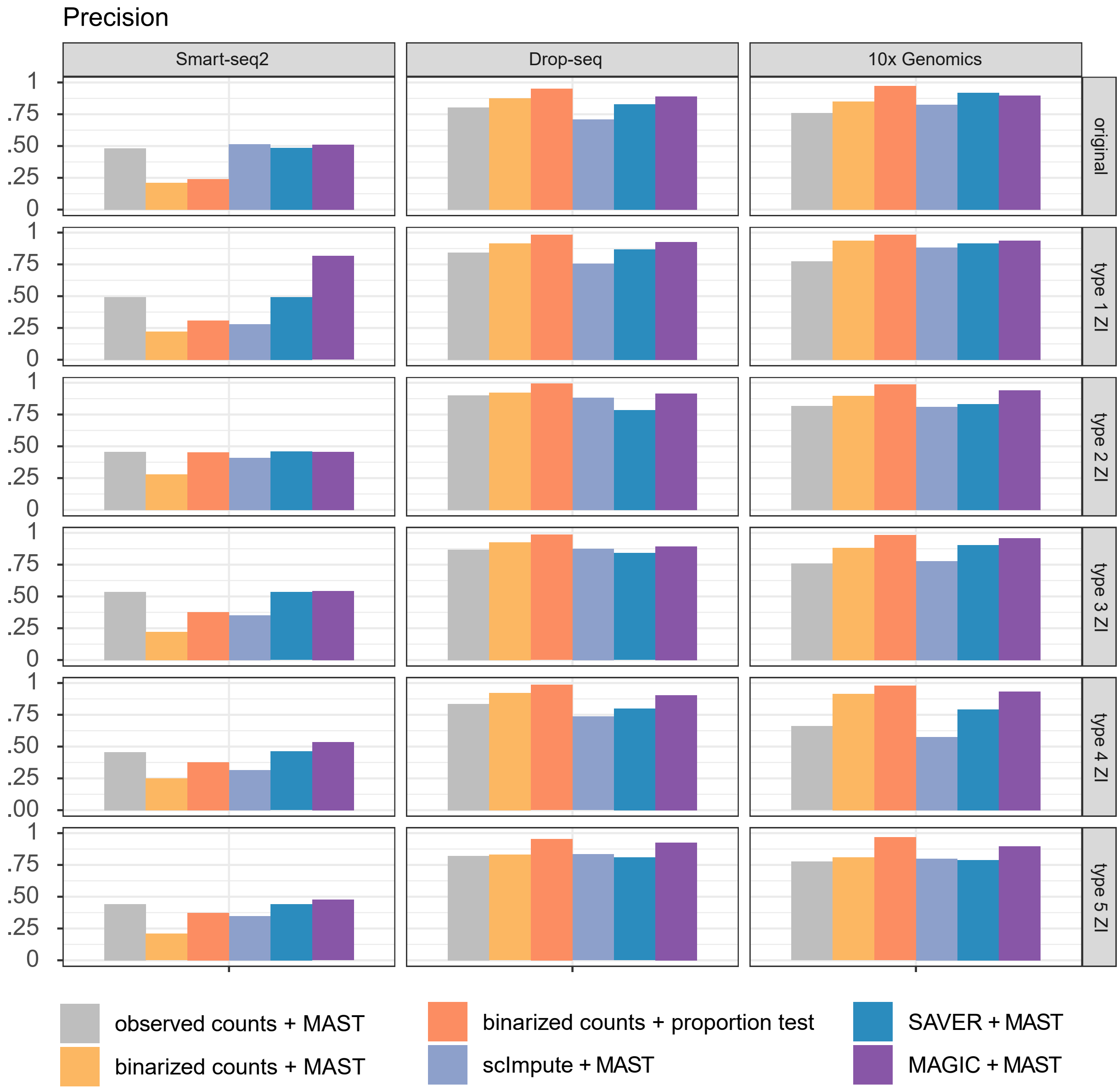


Figure S9: **Evaluation of DE analysis on observed, binarized and imputed data.** We evaluate the DE analysis on Smart-seq2, Drop-seq, and 10x Genomics data based on observed, binarized and imputed data. We perform this analysis before and after using the five masking schemes (type 1 ZI–type 5 ZI) to introduce non-biological zeros. We apply two-sample proportion test on binarzied data and MAST (in Seurat) on observed, binarized, and imputed data to perform DE analysis. We use precision (given the false discovery rate $5\%$) to evaluate the DE results.

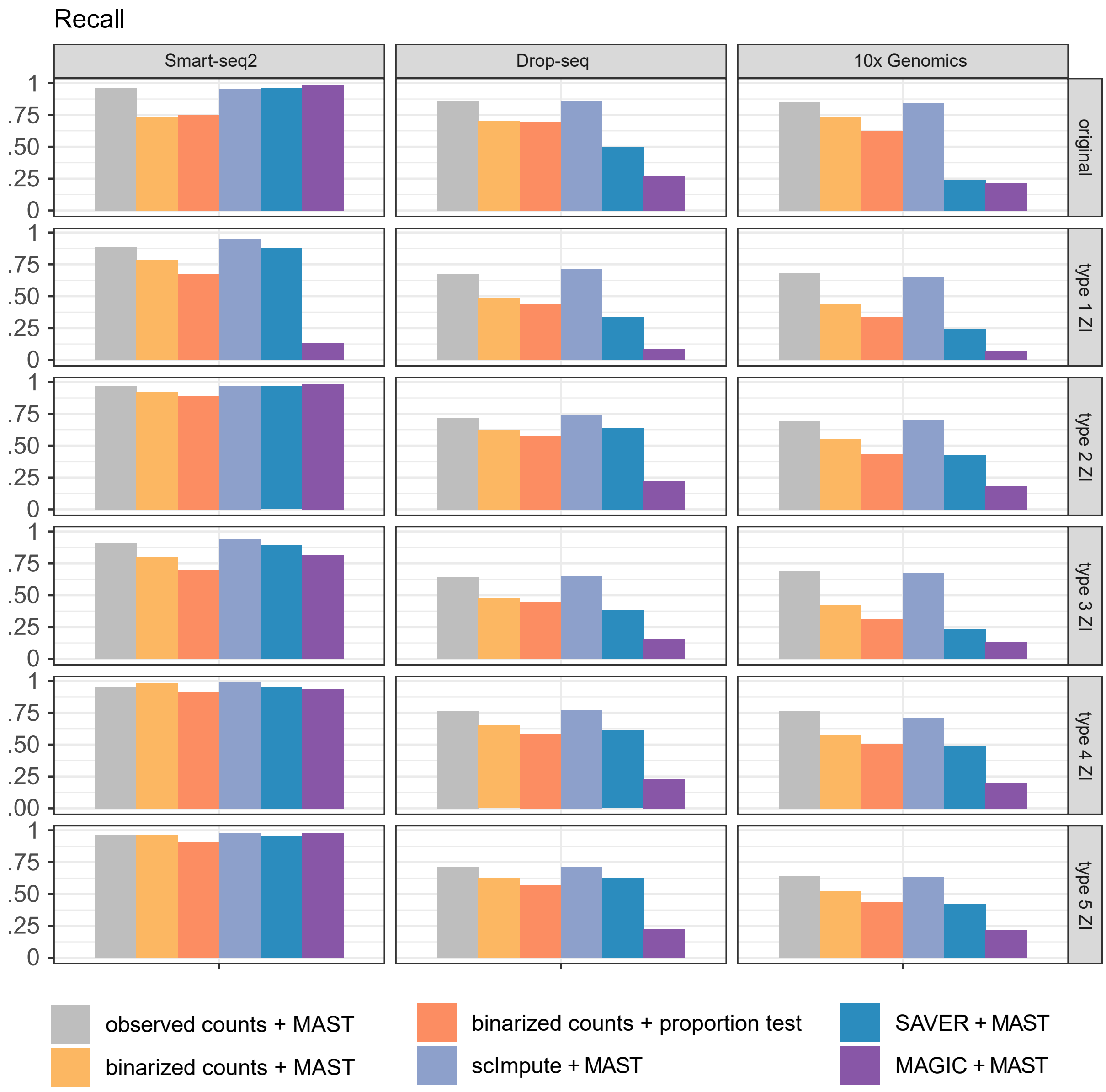


Figure S10: **Evaluation of DE analysis on observed, binarized and imputed data.** We evaluate the DE analysis on Smart-seq2, Drop-seq, and 10x Genomics data based on observed, binarized and imputed data. We perform this analysis before and after using the five masking schemes (type 1 ZI–type 5 ZI) to introduce non-biological zeros. We apply two-sample proportion test on binarzied data and MAST (in Seurat) on observed, binarized, and imputed data to perform DE analysis. We use recall (given the false discovery rate $5\%$) to evaluate the DE results.

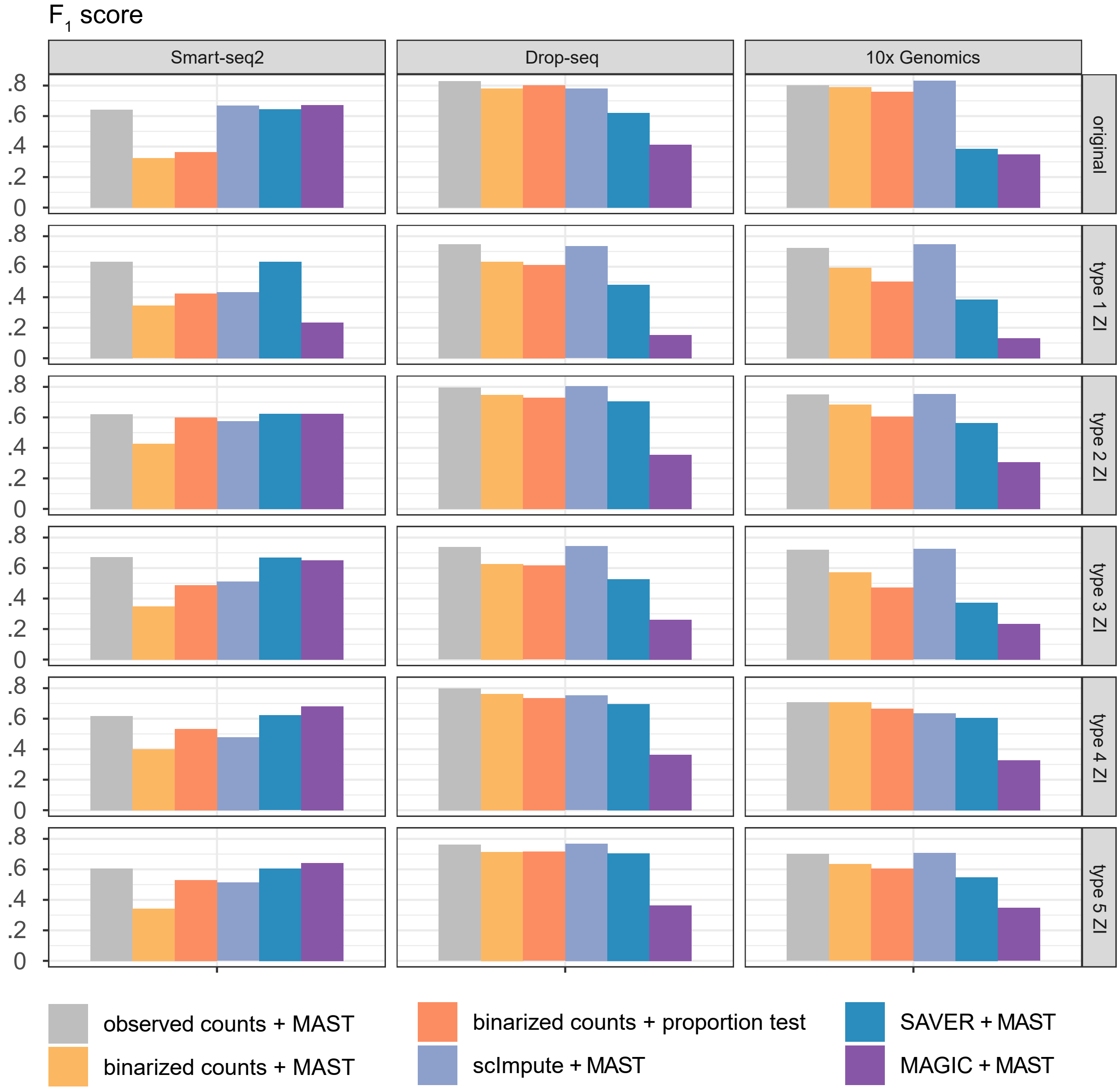


Figure S11: **Evaluation of DE analysis on observed, binarized and imputed data.** We evaluate the DE analysis on Smart-seq2, Drop-seq, and 10x Genomics data based on observed, binarized and imputed data. We perform this analysis before and after using the five masking schemes (type 1 ZI–type 5 ZI) to introduce non-biological zeros. We apply two-sample proportion test on binarzied data and MAST (in Seurat) on observed, binarized, and imputed data to perform DE analysis. We use $F_1$ score (given the false discovery rate $5\%$) to evaluate the DE results.
